# Causally-inspired meta-representation learning framework for predicting patient-specific clinical responses to drug combinations

**DOI:** 10.64898/2026.08.13.744613

**Authors:** Qing-Qing Zhang, Shao-Wu Zhang, Ming-Hui Shi, Jia-Ni Li, Yan-Rui Qiang, Ting-He Zhang

## Abstract

Large-scale prediction and assessment of clinical patient responses (*i.e*., RECIST class) to drug combinations remains challenging due to scarce patient-derived data. The existing prediction methods mainly rely on cancer cell line models. However, substantial biological heterogeneity between cancer cell lines and cancer patients within same tissues, as well as the heterogeneity between one tissue and another, often limit the generalizability of these methods in clinical patients. To overcome these limitations, here we present CaMeRe, a <u>Ca</u>usally-inspired <u>Me</u>ta-representation learning framework designed to predict patient-specific clinical <u>Re</u>sponse to drug combinations. In situations where stable causal factors and domain-specific response-modulating factors are unobservable, explicit discrete domain labels are unavailable, and data is scarce, CaMeRe designed a domain-invariant causal representation learning (DICRL) model guided by the invariant information bottleneck theory and causal intervention invariance principle, and also built a meta-learning framework with bi-level domain generalization to optimize DICRL model for achieving multi-domain generalization within and across tissues. By integrating the causal representation learning and meta learning framework, CaMeRe not only exhibited robust multi-domain generalization performance across multiple clinical drug combination response datasets and PDXs drug combination response datasets and generalization scenarios, but also had better interpretability. We applied CaMeRe to predict drug-combination response scores for 3,423 patients across 542,080 drug combinations. The predicted scores were significantly associated with biomarkers of known drug combinations and enabled the prioritization of candidate drug combinations across 11 cancer types, with stronger support from literature and clinical trial evidences than random baselines. We believe that CaMeRe can be a useful tool for predicting large-scale clinical individual drug combination responses and it has broad clinical applications.

## Introduction

Precision drug combination therapy aims to identify optimal drug sets for individual cancer patients based on their unique omics profiles, thereby overcoming the drug resistance arising from heterogeneity within and between tumors ^1, 2^. Nowadays, the cancer cell lines and *in vitro* models have been extensively applied to drug discovery ^3, 4^. The creation of large-scale pharmacogenomic screening datasets derived from cancer cell lines has significantly advanced our understanding of the intricate relationships among genes, drug combination treatments, and cellular viability. These datasets provide the foundation for training the deep learning models (*e.g*., Deepsynergy ^5^ and MatchMaker ^6^) that predict drug synergy. By leveraging these predictive models, researchers aim to enhance the probability of promising drug combinations successfully passing through clinical trials ^7^. However, owing to our limited understanding of the similarities and differences between cell lines and patient tumors, the drugs with high active in cell line assays may be inactive in some clinical trials, and the response markers identified by cell line experiments may be ineffective in some clinical trials ^8-10^. Even for synergistic drugs that have been tested in clinical trials, the response of cancer patients to these synergistic drugs may be significantly different ^11^. Given this, developing computational methods or models that can illuminate the similarities and differences between cancer cell lines and cancer patients, minimize their discrepancies and capitalize on their commonalities to predict cancer patient responses to drug combination therapies, has emerged as an urgent and critical task in the realm of precision oncology ^12, 13^.

At present, several methods have been developed to predict patient-specific drug responses from preclinical data (*e.g*., cell line data, or *in vitro* screening data), but most of them are designed for single-agent therapies. These preclinical-to-clinical generalization methods can be broadly categorized into the explicit mapping methods and the out-of-domain generalization methods. The explicit mapping methods, such as Celligner ^14^, CELLector ^15^ and HyperTracker ^16^, aim to explicitly match cancer cell lines to the tumor subtypes by evaluating the similarities in phenotypic, genotypic, and gene expression between existing cancer models (*e.g*., cancer cell lines) and the primary tumors. Although these methods can guide the selection of relevant cancer cell lines, they require substantial expert knowledge about genomic alterations in tumor subtypes and the expression profiles of each cell populations (*e.g*., immune cell populations), also assume that the cell lines dataset and tumor datasets have the same subtype composition. Moreover, some studies reveal that many clinical cancer subtypes still lack suitable cell line models ^14-16^, and it is difficult to screen effective drug combination for the matched cell lines from large combination space by trial and error.

The out-of-domain generalization methods mathematically translate the heterogeneity between cancer cell lines and tumors into the data distribution shift, and then design deep transfer learning models to eliminate the distribution shift for predicting drug responses from the cell line pharmacogenomic screening data. According to whether the patient data (*i.e*., target domains) are utilized for model training, out-of-domain generalization methods are divided into the domain adaptive methods (*e.g*., TRANSACT ^17^, CODE-AE ^18^, VAEN ^19^, ADAE ^20^) and domain generalization methods (*e.g*., TCRP ^21^, Velodrome ^22^). Domain adaptive methods utilize the source domain data (*i.e*., cell lines) and target domain data (*i.e*., patients), and employ the domain-adversarial learning or maximum mean discrepancy approaches to align the distribution of cell lines and clinical patients either in the input/feature space, or output space for predicting drug responses of cancer patients. However, patient data is usually unavailable in drug response prediction task, because we do not know who will become cancer patients. Domain generalization methods aim to generalize a model trained with multiple source domains to the unseen test domains (*i.e*., target domains) ^23, 24^. For example, Velodrome ^22^ takes the cell lines or unlabeled patients as source domain data to learn domain-invariant features and hypothesis-invariant features, and then predicts drug response of other patients (*i.e*., unseen target domain) by integrating multiple domain-specific predictors. TCRP ^21^ takes the cell lines or PDX models from different tissues as source domain data to learn domain-invariant meta-knowledge (*i.e*., the initial parameters of TCRP) via meta-learning, and then predicts drug response of cell lines or patients in other tissues (*i.e*., unseen target domain) based on the learned meta-knowledge. However, existing domain generalization methods generally focus on modeling the dependency between domain-invariant transcriptome features and drug response, while overlooking the intrinsic causal mechanisms through which drug response is jointly determined by transcriptomic and drug or drug-combinations features. As a consequence, such models may generalize poorly when causal features presumed to be domain-invariant differ across drug or drug combinations, or when non-causal but apparently domain-invariant features shift in unseen target domains.

In the representation learning stage, several causality-aware domain generalization methods have successfully leveraged different causal theories to learn representations that capture underlying domain-invariant causal mechanisms, thereby improving their generalization performance ^23, 25, 26^. In addition, considering that preclinical drug screening is fundamentally predicated on the existence of partially shared drug-response mechanisms between preclinical models and patient, in this work we will introduce the causal invariance theory into our domain generalization framework for predicting cancer patient responses to drug combinations. Specifically, from a structured causal mechanistic perspective, we aim to accentuate the shared drug combination response-relevant mechanisms that are relatively stable and transferable across cell lines and patients, while attenuating domain-dependent pathways that drive unstable or non-transferable associations between input (*i.e*., the joint features of drug A, drug B, and a cell line/patient) and output (*i.e*., response to drug combination), thereby achieve better generalization ability on predicting patient-specific response to the drug combination. However, due to the complex nature of the combined features of drug A, drug B, and a cell line/patient, directly adopting existing causality aware domain generalization methods faces the following challenges. First, causality aware domain generalization methods developed on other fields (*e.g*., computer vision) often rely on explicit environment (or domain) labels for distinguishing the sources of distribution shift, whereas such labels are expensive to obtain and often unavailable for the joint features of drug A, drug B, and a cell line/patient. Second, the causality aware domain generalization methods are often designed for specific task, requiring sufficient and diverse domains to enhance the generalization, while the number of cell lines or patients combined with specific drug combination in specific tissue is small, even zero. Therefore, it is challenging to train a domain generalization model that is robust to domain shift between cell lines and patients for a specific drug combination.

To tackle these challenges, we will fully utilize the advantages of meta-learning and causality aware domain generalization methods, thus propose a <u>Ca</u>usally-inspired <u>Me</u>ta-representation learning framework (named CaMeRe) to predict patient-specific clinical <u>Re</u>sponses to drug combinations by capturing the stable and transferable action mechanism of each specific drug combination in specific tissue, because distilling the experience (*e.g*., good weight initialization) learned from multiple tasks (*e.g*., source tissues) by meta-learning process can learn a new task (*e.g*., target tissue) fast with a few training samples. CaMeRe first establishes a structural causal model (SCM) to describe potential distribution shifts in drug-combination response prediction, where predictions are made from the joint representation of drug-combination features and transcriptomic features of cell lines or patients. Then, under the SCM-based causal assumption, based on the invariant information bottleneck theory ^27, 28^ and causal intervention invariance principle ^29^, CaMeRe designs five tractable loss functions for the inner-loop and outer-loop optimization of meta-learning process to eliminate distribution shifts among cell line samples and patient samples in specific tissues, as well as distribution shifts among different tissues, which enables CaMeRe to extract the causal latent representation from the low-level input features for predicting unseen clinical patient response to drug combinations. Due to the inability to obtain explicit discrete domain labels, CaMeRe presumes that causal latent representation and domain-specific latent representation belong to multivariate diagonal gaussian distribution, thus designs an unsupervised multi-domain discriminator to define the unidentifiability of causal latent representation and the identifiability of domain-specific latent representation according to the exponential of Wasserstein distance. Owing to the reconstruction of the invariant causal mechanism between combination features and combination response, CaMeRe can identify the important genes through post-hoc interpretability models (*e.g*., SHapley Additive exPlanations) and has capability to assist causal inference.

To evaluate the effectiveness of CaMeRe, we performed extensive comparative studies against several existing methods, and the results show that CaMeRe can effectively predict drug combination response of clinical patients across diverse tissue types (such as breast, lung, ovary, and colon), and also identify genes with significant causal effects on the response to a specific drug combination. In addition, we applied CaMeRe to predict the response scores of 3,423 cancer patients (covering 11 types of cancer) to 5.4 million drug combinations, and prioritized several candidate drug combinations that are significantly enriched in high-response groups of each cancer, some of which have already been reported in existing clinical trials and literatures.

## Results

### A. Overview of CaMeRe

CaMeRe is a causally-inspired meta representation learning framework designed to transfer drug-combination response knowledge from cell lines to patients, enabling large-scale prediction of patient responses to drug combinations, prioritization of candidate drug combinations, and nomination of candidate causal genes for downstream causal effect estimation. An overview of CaMeRe framework is illustrated in **Fig.1**, and detailed methodological descriptions are provided in the Methods section.

**Fig 1.**
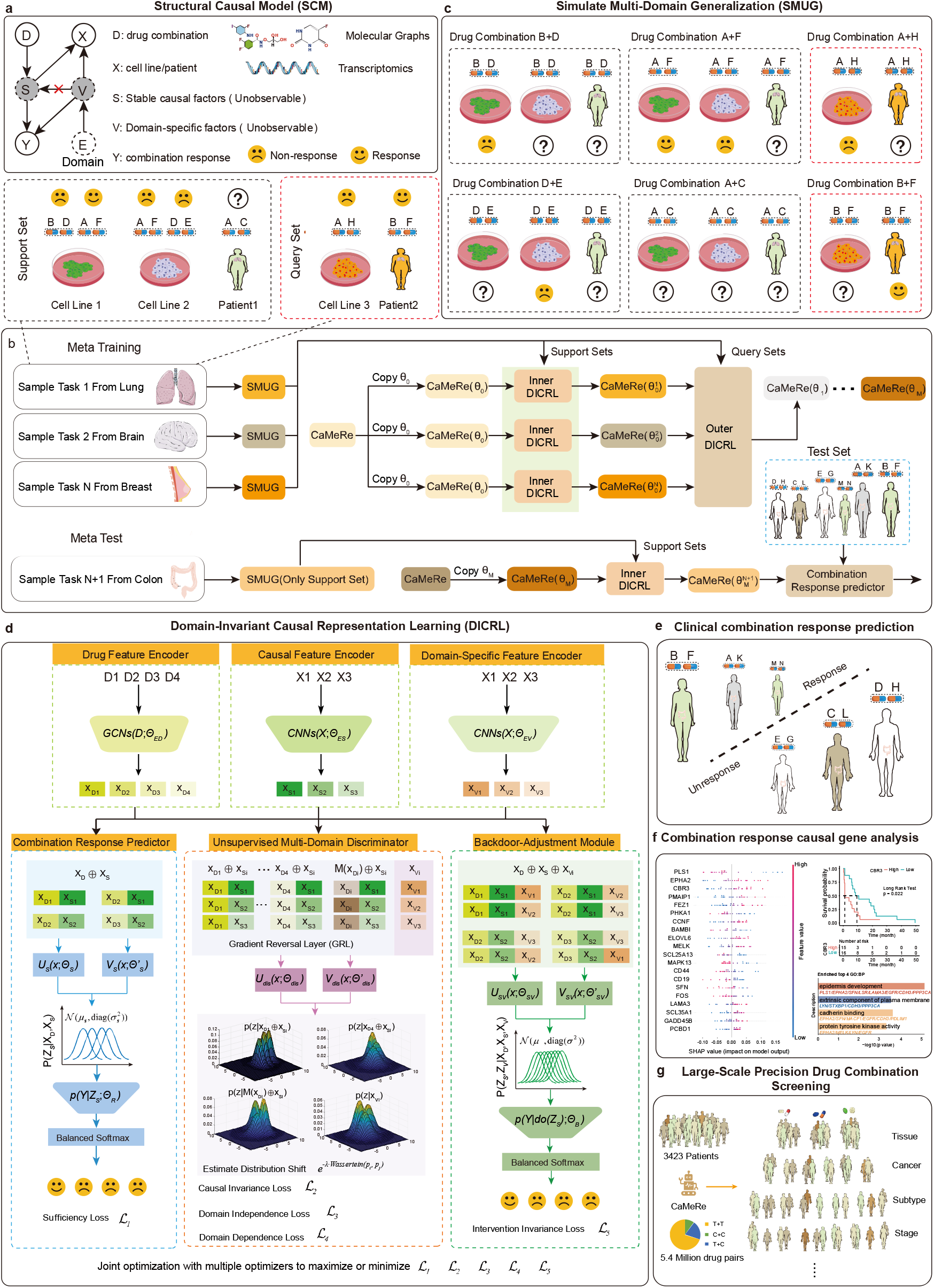
Overview of CaMeRe, a causally-inspired meta representation learning framework for predicting patient-specific clinical drug combination response. **a**, The Structure Causal Model (SCM) for formalizing the framework of predicting clinical drug combination response. **b**, Meta-Training stage and Meta-test stage. **c**, Simulating multi-domain generalization scenario based on the support set and query set of each sampled task. For each drug combination, each cell line or patient belongs to a domain, where 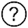 donates that the label Y is unknown. **d**, Taking the support set of Task 1 as an example for illustrating the domain-invariant causal representation learning (DICRL) process. DICRL aims to optimizing CaMeRe by minimize the sufficiency loss, causal invariance loss, domain independence loss, domain dependency loss, and intervention invariance loss. Specifically, the inner DICRL utilizes each support set to optimize CaMeRe by minimize the five losses for each task, while the outer DICRL utilizes the query sets of a batch of tasks to optimize CaMeRe by minimize the five mean losses for the batch tasks. **e**, Predicting clinical patients’ response to different drug combinations with the trained CaMeRe. **f**, Illuminating the ‘causal’ genes of patient responses to specific drug combination based on SHAP important scores. **g**. Screening the potential drug combination from 5.4 million drug pairs for 3,423 TCGA patients.

The input of CaMeRe can be any combined features consisting of two drug molecular graphs and a cell line/patient’s transcriptomic features, and the output is the drug combination response of the cell line/patient. Patients with complete response (CR) or partial response (PR) are defined as responders, whereas those with stable disease (SD) or progressive disease (PD) are defined as non-responders according to RECIST criteria. CaMeRe first takes the drug combination response samples in each cell line or patient as a domain and constructs a structural causal model (**Fig.1a**) to characterize the intrinsic causal relationship between the combined input features and drug-combination response, thereby improving the generalization ability of CaMeRe. Then, CaMeRe performs meta-learning at the level of tissue-specific prediction tasks (**Fig.1b**), and each task simulates multi-domain shift across cell lines and patients in specific tissue (**Fig.1c**) to eliminate the inherent bi-level biological heterogeneity within and between tissues.

The causally inspired meta-representation learning process (**Fig.1b**) in CaMeRe comprises four main phases: multi-domain shift simulation (**Fig.1c**), inner/outer domain invariant causal representation learning (**Fig.1d**), and clinical test (**Fig.1e**). For each task, CaMeRe builds a support set and a query set to simulate multi-domain shift in a tissue (**Fig.1c**). The support set consists of the labeled samples of few cell lines and the unlabeled samples of few patients, and the query set consists of the labeled samples of other few patients and the labeled samples of other few cell lines in the tissue. The cell lines or patients in support set and query set do not overlap. Due to very small number of the labeled patient samples, CaMeRe takes the labeled samples of other few cell lines as auxiliary domains to reduce overfitting.

Due to the unobservability of stable causal factors and domain-specific response-modulating factors hidden in the combined features, CaMeRe instead extracts the causal/domain-specific latent representations from the combined features, where these latent representations have the same attributes as stable causal factors and domain-specific response-modulating factors, and then builds the invariant causal mechanisms with the aid of causal intervention. Under the SCM assumptions and guided by the invariant information bottleneck theory and causal intervention invariance principle, CaMeRe operationalizes the above theoretical objectives as five tractable loss functions: sufficiency loss, causal invariance loss, domain independence loss, domain dependence loss, and intervention invariance loss, to ensure the causal latent representation specific to each drug combination, independent on cell lines or patients, having all stable and transferable information about drug combination response, while the domain-specific latent representation dependent on cell lines or patients, without the stable and transferable information about drug combination response. Specifically, the sufficiency loss function is used to measure the difference between the true drug combination response and the predicted drug combination response based on causal latent representation; the causal invariance loss function is used to measure the causal latent distribution shift between cell lines and patients for specific drug combination; the domain independence loss function is used to measure the drug-averaged causal latent distribution shift between cell lines and patients; the domain dependence loss function is used to distinguish domain-specific latent distribution between cell lines and patients; the causal intervention invariance loss function is used to measure the difference between the true drug combination response and the predicted drug combination response based on the intervening causal latent representation.

In inner domain invariant causal representation phase, CaMeRe fine tunes on each task support set to simultaneously minimize five loss functions. In outer domain invariant causal representation phase, CaMeRe takes cell lines and patients in query set as multiple unseen domains to test the power of the fine-tuned CaMeRe for a specific task. By updating the CaMeRe with the mean loss of a batch of tasks, the updated CaMeRe can easily adapt new tissue scenarios.

In clinical test phase, the trained CaMeRe is transferred to new tissue, and the support set of new tissue is used to fine-tune CaMeRe by inner domain invariant causal learning process, thus the fine-tuned CaMeRe is used to perform the downstream tasks, such as predicting patient response to any drug combination, analyzing the important genes of combination response through post-hoc interpretability models, and screening potential anti-cancer drug combinations at large scale.

### B. Results of CaMeRe and other comparative methods on clinical patients

To evaluate the performance of CaMeRe in predicting clinical drug combination response, we retrieve the clinical drug combination response data from The Cancer Genome Atlas (TCGA) ^30^, and the cancer cell line drug synergy data from DrugComb ^31, 32^. Based on tumor primary site, we group the clinical patients and cancer cell lines into five tissues, including Breast tissue (TP =86, TN=10, NUP= 238, NCL=11), Colon tissue (TP=38, TN=26, NUP= 29, NCL=11), Ovary tissue (TP=51, TN=16, NUP= 352, NCL=12), Lung tissue (TP=90, TN=32, NUP= 66, NCL=13), and Brain tissue (TP=2, TN=13, NUP= 29, NCL=6). Here, TP, TN, NUP, and NCL denote the number of true positive samples, the number of true negative samples, the number of unlabeled patient samples, the number of cancer cell lines, respectively. Each of the 5 tissue-specific clinical drug combination response dataset is used in turn as the meta-testing dataset, while the remaining four tissue-specific datasets are used as meta-training dataset. Notably, for a meta-testing dataset, only the cell lines samples and unlabeled patient samples in the meta-testing tissue are used to fine-tune the pretrained model.

For the new task of clinical drug combination response prediction, we first improve some previous methods as baseline methods to predict the clinical drug combination response. These methods that need to be modified mainly include the feature engineering-based methods for predicting drug synergy (*i.e*., XGBoost, DeepSynergy, Matchmaker), and the domain adaptive-based methods for predicting clinical drug response (*i.e*., ADAE, DSN-M, DSN-A, CODE-AE), and the domain generalization methods (*i.e*., Velodrome, TCRP) for predicting clinical drug response. These modified methods are named as XGBoost^+^, DeepSynergy^+^, Matchmaker^+^, ADAE^+^, DSNM^+^, DSNA^+^, CODE-AE^+^, Velodrome^+^, TCRP^+^(see Supplementary Information **Note 1** for details). Then, we systematically compare the performance of our CaMeRe against these baseline methods by adopting the metrics of Area Under the Receiver Operating Characteristic Curve (AUROC), Area Under the Precision-Recall Curve (AUPR), Balanced Accuracy (BACC), and Matthews Correlation Coefficient (MCC). To ensure a fair comparison, all methods are trained and tested using the same data splitting, repeating five times, and the average results along with standard deviation over five repeated runs for each of the five tissues is calculated. In addition, to increase statistical power, the overall mean ± standard deviation is calculated by combining the results from all five tissues across the five runs. These results are shown in **Fig.2, Fig.S1**, and **Tables S1-S5** (in Supplementary Information). When statistically comparing one method with other methods, p-values are adjusted using the multiple comparison correction (Benjamini-Hochberg method) procedure to control the false discovery rate.

**Fig 2.**
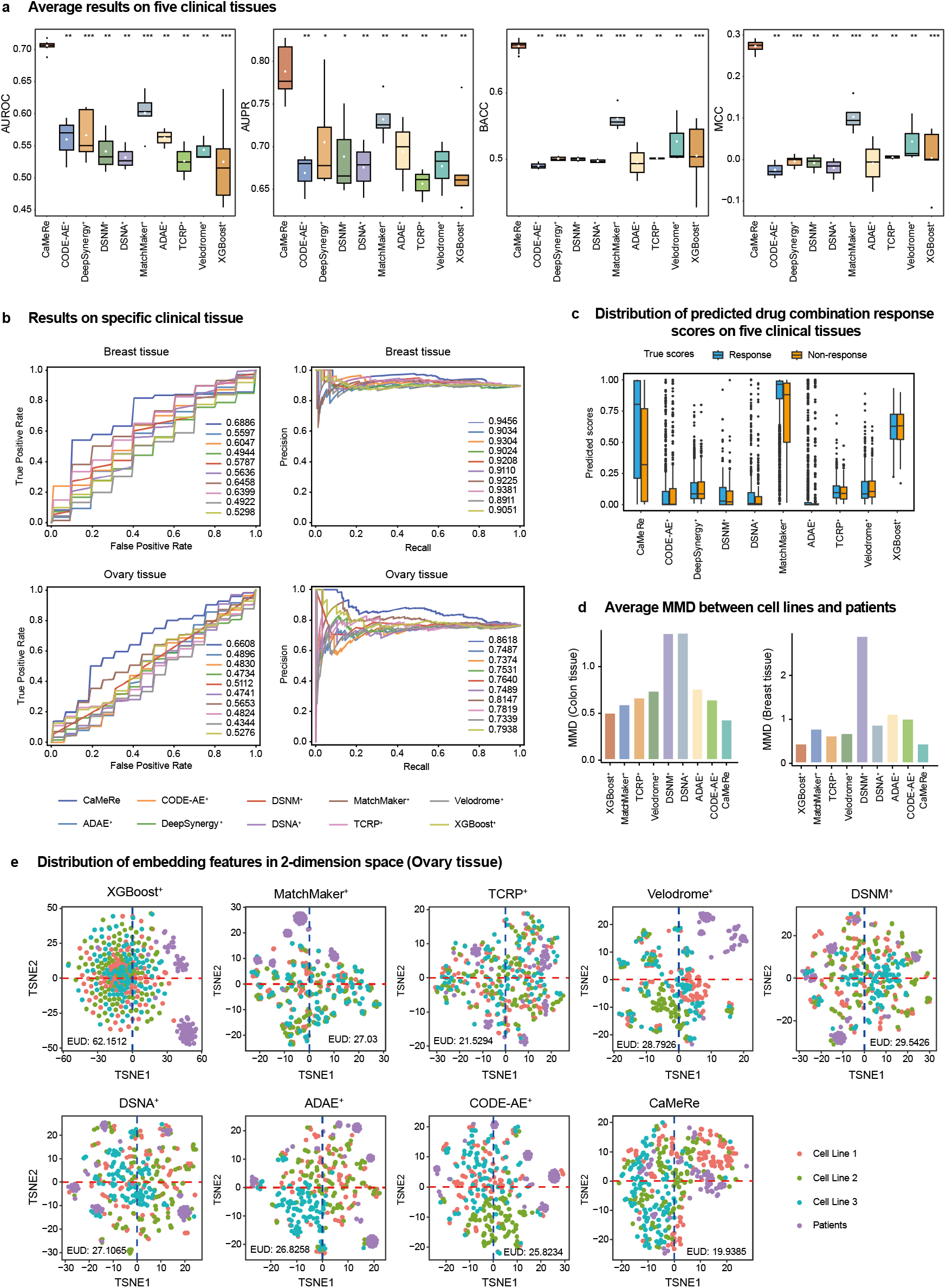
Results of CaMeRe and other comparative methods on five clinical tissue datasets. **a**, Box plots of average AUROC, AUPR, BACC, and MCC over all clinical tissue datasets, with each method repeated five times. The center line in the box represents the median among the five experimental results, excluding outliers; the lower and upper bounds of the box represent the first (Q1) and third (Q3) quartiles, respectively; the lower and upper bounds of the whiskers represent the minima and maxima, excluding outliers, respectively; the white dot in the box represents the mean among the five experimental results. The two-sided Wilcoxon signed-rank test is selected to compute the significant difference between CaMeRe and other baselines (ns denotes p>0.05, * denotes p<0.05, ** denotes p<0.01, *** denotes p<0.005, **** denotes p<0.001). **b**, Receiver Operating Characteristic curve and Precision-Recall curve on Breast tissue dataset and Ovary tissue dataset, respectively. **c**, Box plots of the predicted combination response scores by CaMeRe and other comparative methods on the five clinical tissue datasets. The predicted combination response scores are grouped by the ground truth (*i.e*., response or non-response). **d**, Bar plots of the average Maximum Mean Discrepancy (MMD) between three cell lines domains in support set and patient domains in query set from Colon tissue dataset and Breast tissue dataset. The MMD is calculated based on the feature representations learned by CaMeRe and other baselines. **e**, T-SNE visualization of the feature representations learned by CaMeRe and other methods on Ovary tissue dataset. Here, “EUD” denotes the EUclidean Distance between feature representations in two-dimensional embedding space.

As shown in **Fig.2, Fig.S1**, and **Tables S1-S5**, our CaMeRe outperforms all baseline methods in predicting the clinical drug combination responses, both on each tissue-specific tissue (*i.e*., Ovary, Breast, Colon, Lung, and Brain) and the overall average across all tissues. Despite severe class imbalance and significant distribution shifts in the dataset, the average AUROC, AUPR, BACC, and MCC of our CaMeRe are 0.71, 0.79, 0.67, and 0.27, respectively, while all the comparison methods have AUROC less than 0.65, AUPR less than 0.75, BACC less than 0.60, and MCC less than 0.15. Specifically, in the overall average across all tissues, compared to MatchMaker^+^ method that has the best performance in baseline methods, the AUROC, AUPR, and BACC of our CaMeRe increases by 24.1%, 18.4%, and 21.3%, respectively. Moreover, CaMeRe has also achieved the significant performance improvements in specific-tissue scenarios. For example, the AUROC, AUPR, and BACC of our CaMeRe for Ovarian tissue are 9.7%, 2.2%, and 23.4% higher than that of MatchMaker^+^, respectively; for Breast tissue, the AUROC, AUPR, and BACC of our CaMeRe are 7.6%, 0.8%, and 21.1% higher than that of MatchMaker^+^, respectively. These results demonstrate that even on the clinical data with severe class imbalance and domain data distribution shifts, our CaMeRe can effectively predict drug combination response of patients. Additionally, we also noted that the domain adaption and domain generalization methods, such as ADAE^+^, CODE-AE^+^, TCRP^+^, DSNM^+^, DSNA^+^, Velodrome^+^, cannot effectively distinguish between positive and negative samples (**Fig.2c** and **Fig.S2**). Specifically, these baseline methods consistently achieve prediction scores around 0.1 for all samples, indicating that they are unable to mitigate the impact of label bias arose from severe data imbalance.

Besides, to qualitatively evaluate the capacity of our CaMeRe for addressing domain shift between cell-line samples and patient samples, we applied *t*-distributed Stochastic Neighbor Embedding (t-SNE) to projected the learned representations from CaMeRe and other comparison methods into two dimensions for visualization. As illustrated in **Fig.2e** and **Fig.S3**, we can see that CaMeRe effectively reduces the spatial separation between patient samples and cell line samples. Moreover, we quantitatively assess the alignment between patient and cell-line distributions using Maximum Mean Discrepancy (MMD). As shown in **Fig.2d** and **Fig.S4**, the average MMD of CaMeRe is the smallest.

These results demonstrate that our CaMeRe is significantly more effective than other domain adaptation and generalization methods in mitigating domain shifts, thereby improving the generalizability of CaMeRe in predicting clinical drug combination response.

### C. Results of CaMeRe and other comparative methods on PDXs

To further evaluate the performance of CaMeRe in predicting combination response of Patient-Derived Xenografts (PDXs), we retrieve PDXs drug combination response data from previous paper ^33^, unlabeled clinical drug combination response data from TCGA, and cancer cell line drug synergy data from DrugComb ^31, 32^. Subsequently, based on tumor tissue or tumor primary site, we group the PDXs, clinical patients, and cell lines into four tissues, that is colon tissue (TP = 22, TN = 161, NUP= 89, NCL=11), skin tissue (TP = 17, TN = 96, NUP= 12, NCL=14), breast tissue (TP = 21, TN = 54, NUP= 328, NCL=11), and lung tissue (TP = 13, TN = 62, NUP= 184, NCL=13). Then, each of the 4 tissue-specific PDXs drug combination response dataset is used in turn as the meta-testing dataset, while the remaining three tissue-specific datasets are used as meta-training dataset. For a meta-testing dataset, only the cell lines samples and unlabeled patient samples in the meta-testing tissue are used to fine-tune the model.

We test the performance of CaMeRe and other nine methods (*i.e*., XGBoost^+^, DeepSynergy^+^, Matchmaker^+^, TCRP^+^, CODE-AE^+^, Velodrome^+^, ADAE^+^, DSNM^+^, DSNA^+^) on the drug combination response datasets of Patient-Derived Xenograft. To ensure the fairness in comparative analysis, we repeat five times for each method when each tissue-specific dataset is used as meta-testing dataset. The results of CaMeRe and other nine methods are shown in **Fig.3, Fig.S5**, and **Tables S6-S9**.

**Fig 3.**
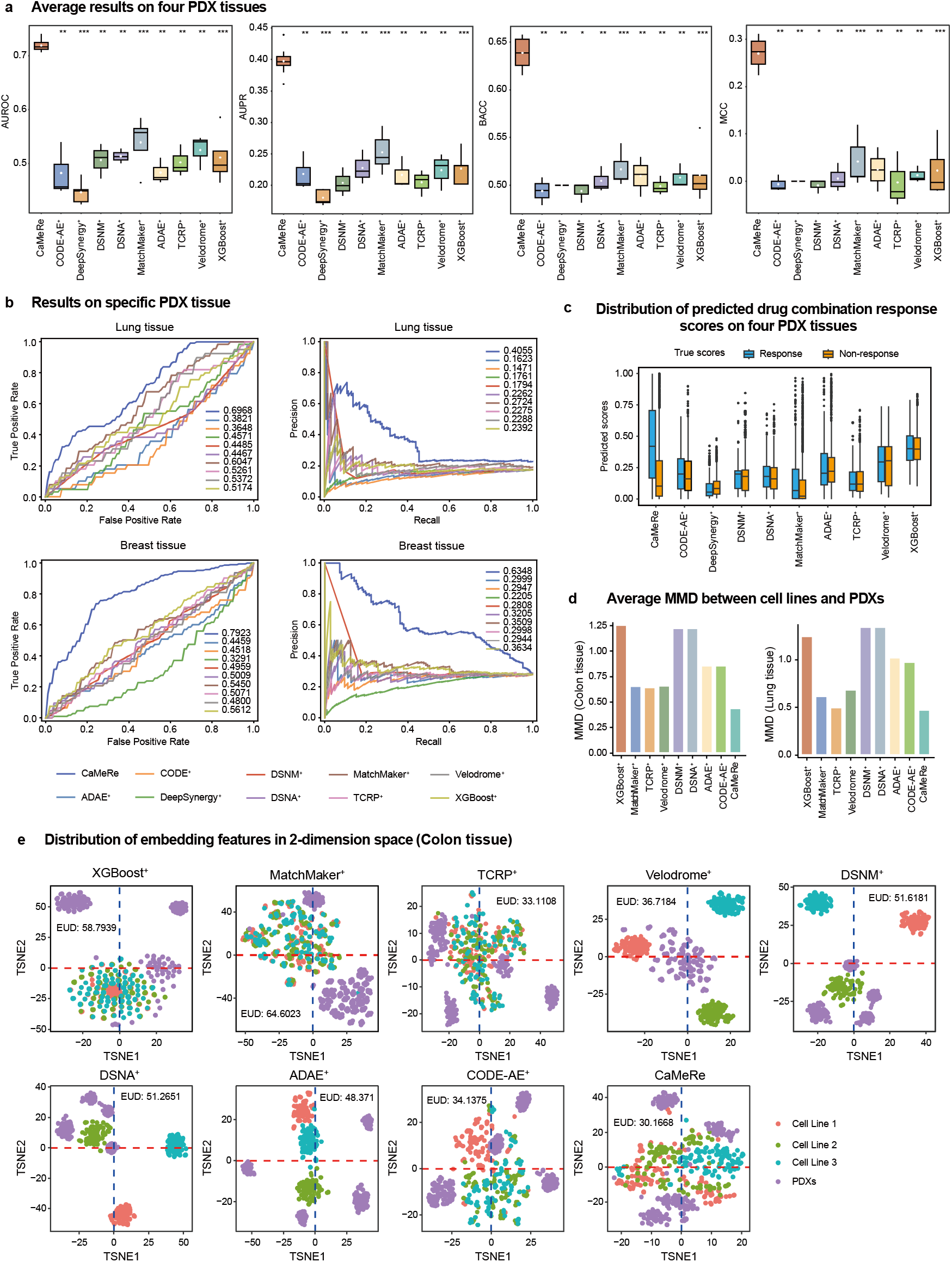
Results of CaMeRe and other comparative methods on four PDX tissues. **a**, Box plots of average AUROC, AUPR, BACC, and MCC over four PDX tissues, with each method repeated five times. The two-sided Wilcoxon signed-rank test is used to compute the significant difference between CaMeRe and other comparative methods. **b**, Receiver Operating Characteristic curve and Precision-Recall curve on Breast tissue dataset and Lung tissue dataset. **c**, Box plots of the predicted combination response scores by CaMeRe and other comparative methods on the four PDX tissue datasets. The predicted combination response scores are grouped by the ground truth (*i.e*., response or non-response). **d**, Bar plots of average Maximum Mean Discrepancy (MMD) between three cell lines domains in support set and PDX domains in query set from Colon tissue and Lung tissue. **e**, T-SNE visualization of the feature representations learned by CaMeRe and other comparison methods on Colon tissue.

From **Fig.3, Fig.S5**, and **Tables S6–S9**, we can see that our CaMeRe consistently surpasses all comparison methods, achieving superior results on each specific tissue (*i.e*., Colon, Skin, Breast, or Lung) as well as the overall average across all tissues. In term of the overall average across all tissues, the average AUROC, AUPR, BACC and MCC of CaMeRe are 24.1%, 18.4%, 21.3%, and 258.5% higher than the best performing MatchMaker^+^ among nine comparison methods. Simultaneously, compared to the best baseline method, the AUROC, AUPR, BACC and MCC of CaMeRe increases by 15.2% and 48.9%, 17.9%, and 260.0% for Colon cancer, and 41.2%, 74.7%, 31.8%, and 412.1% for Lung cancer, respectively. From **Fig.3c** and **Fig.S6**, we can see that on the PDX drug combination response datasets with a higher proportion of non-response samples, the prediction scores (median) of CaMeRe and MatchMaker^+^ for responders are 0.42 and 0.07, respectively, while for non-responders they are 0.1 and 0.02, respectively, meaning that MatchMaker^+^ wrongly predicts more responders as non-responders. From **Fig.2c**, we can see that on the clinical drug combination dataset with a higher proportion of response samples, the prediction scores (median) of CaMeRe and MatchMaker^+^ for responders are 0.8 and 0.96, respectively, while for non-responders they are 0.32 and 0.88, respectively, meaning that MatchMaker^+^ wrongly predicts more non-responders as responders. These results indicate that for the datasets with imbalanced responder and non-responder samples, MatchMaker^+^ favors the category with more samples, resulting in higher false positive/false negative rates, while our CaMeRe can effectively mitigate the impact of label bias caused by sample label imbalance. In addition, from **Figs.3d-3e, Figs.S4** and **S7**, we can see that the feature representations of three cell-line sample groups and PDX samples learned by CaMeRe exhibited the minimal distributional gap among all methods, indicating CaMeRe superior effectiveness in mitigating domain shift compared to other domain generalization methods. Moreover, CaMeRe effectively preserves the spatial specificity of drug combinations in the embedding space.

Above results demonstrate that our CaMeRe is significantly more effective than baseline methods in terms of generalizability from cancer cell lines and unlabeled patients to PDX models on a specific tissue, from one tissue to another tissue.

### D. Ablation Studies

To evaluate the contribution of each key component in CaMeRe, we design 8 variants of CaMeRe by individually removing one key component in CaMeRe. **Figs.4a-4b** display the average results of CaMeRe ablation experiments on five tissue-specific clinical drug combinations response datasets (*i.e*., Ovary, Breast, Colon, Lung, and Brain), as well as the average results of CaMeRe ablation experiments on four tissue-specific PDXs drug combinations response datasets (*i.e*., Colon, Skin, Breast, and Lung), respectively. **Figs.S8-S9** display the ablation results of CaMeRe on each tissue-specific clinical drug combinations response dataset and PDX drug combinations response dataset, respectively. Here, **CaMeRe w/o BA** denotes the removal of the backdoor-adjustment block from CaMeRe framework. **CaMeRe w/o IDS** denotes the removal of the constraint on inner distribution shift. **CaMeRe w/o DNC** denotes the removal of the constraint distinguishing domain-specific latent representation. **CaMeRe w/o DCS** denotes the removal of the constraint distinguishing causal latent representation unique to specific drug combinations. **CaMeRe w/o DCA** denotes the removal of the constraint distinguishing causal latent representation shared across arbitrary drug combinations. **CaMeRe w/o LSD** denotes the removal of the latent space distribution constraint. **CaMeRe w/o AT** denotes the removal of the auxiliary task constraint, implying that CaMeRe excludes combination response samples of cell lines from each query set. **CaMeRe w/o BS** denotes the replacement of the balanced softmax loss with standard softmax loss.

**Fig 4.**
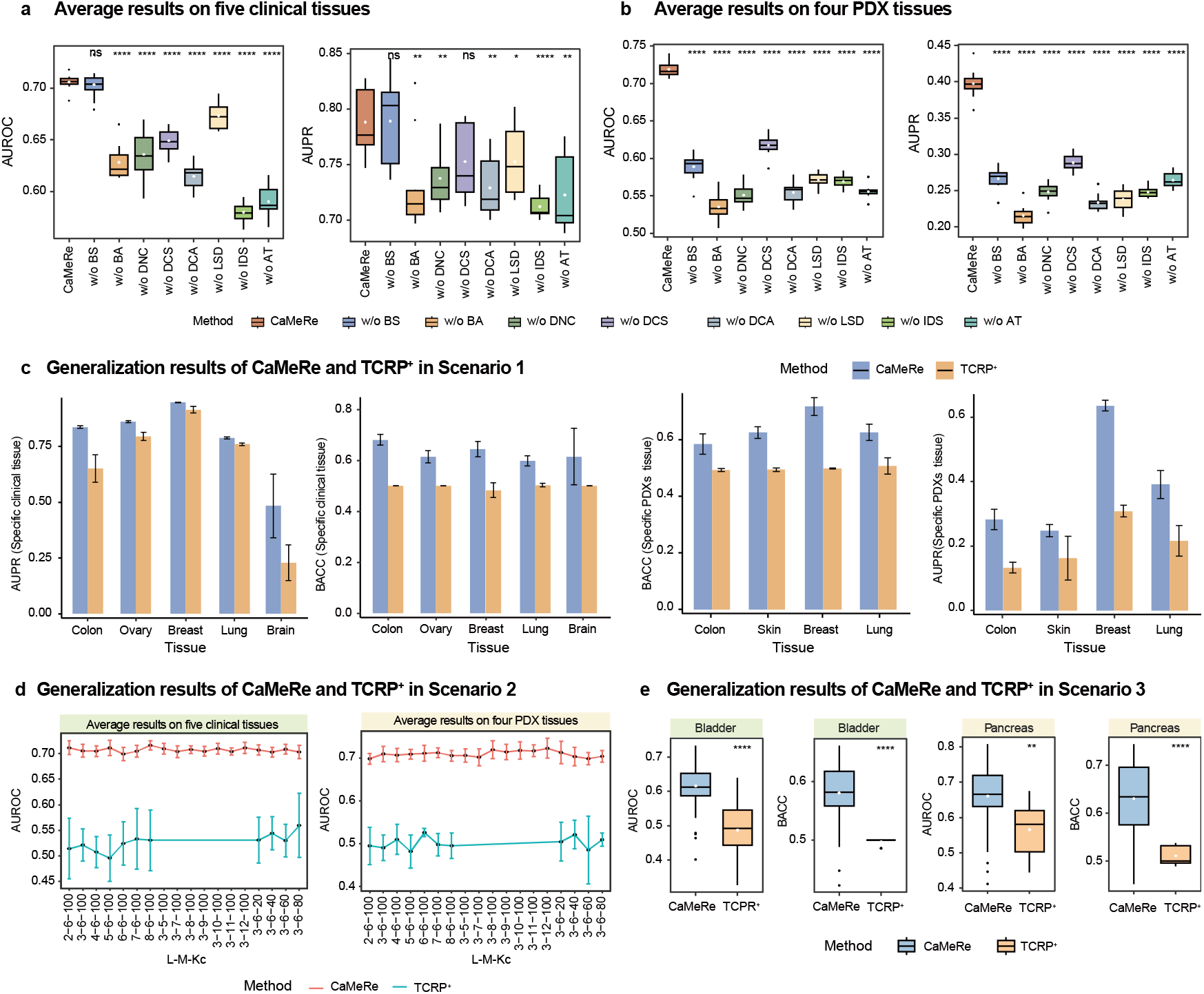
Ablation results of CaMeRe, and the generalization results of CaMeRe and TCRP^+^. **a-b**, Box plots of average AUROC and AUPR across all clinical tissue datasets (**a**) and all PDX tissue datasets (**b**), with each method repeated five times. The two-sided Wilcoxon signed-rank test is used to compute the significant difference between CaMeRe and its variants. **c**, Bar plots of AUPR and BACC generated by CaMeRe and TCRP^+^ on five clinical tissue datasets and four PDX tissue datasets in the first domain generalization scenario, in which error bars represent the standard deviation across five independent runs. **d**, Line graphs of the average AUROC of CaMeRe and TCRP^+^ on five clinical tissue datasets and four PDX tissue datasets in the second domain generalization scenario, in which error bars represent standard deviation across five independent runs. **e**, Box plots of the average AUROC and BACC of CaMeRe and TCRP^+^ on Bladder tissue dataset and Pancreatic tissue dataset in the third domain generalization scenario.

From **Figs.4a-4b**, we observe that the performance of CaMeRe w/o IDS is the worst on clinical drug combination response datasets. Specifically, the average AUROC, AUPR, BACC, and MCC of CaMeRe w/o IDS on clinical drug combination response datasets decrease by 17.8%, 9.6%, 14.6%, and 56.0%, respectively. These results underscore the critical role of eliminating the distribution shift between support set (*i.e*., the cell lines samples and unlabeled patient samples) and query set (*i.e*., the labeled patient samples) in each specific tissue. We also observe that the performance of CaMeRe w/o BA is the worst on PDXs drug combination response datasets. Specifically, the average AUROC, AUPR, BACC, and MCC of CaMeRe w/o BA on PDXs drug combination response datasets decrease by 25.5%, 46.0%, 19.4%, and 93.3%, respectively. The most severe performance degradation highlights the importance of removing the confounding effects introduced by domain-specific latent representation, particularly in the context of domain generalization.

In addition, from **Figs.4a-4b**, we find that the average AUROC, AUPR, BACC and MCC of CaMeRe w/o DNC, CaMePa w/o LSD, CaMePa w/o DCS, and CaMePa w/o DCA are consistently lower than that of CaMeRe both on the clinical drug combination response datasets and the PDXs drug combination response datasets. These results indicate that enforcing causal latent representation and domain-specific latent representation to share similar attributes with stable causal factors and domain-specific factors are more desired for CaMeRe.

From the results of CaMeRe w/o AT and CaMeRe w/o BS in **Figs.4a-4b**, we find that removing each of two training tracks (*i.e*., adding auxiliary task, balancing label distribution) will downgrade the prediction performance of CaMeRe. The reason is that the labeled patient samples and the labeled PDX samples are very small, adding the labeled cell line samples as auxiliary task can reduce the overfitting during meta-training. Moreover, the imbalance between positive samples and negative samples leads to the label distribution shift, adding the constraint on balancing sample distribution can reduce the label shift and improve the prediction performance.

Overall, the results of **Figs.4a-4b** and **Figs.S8-S9** show that removing any key component will seriously affect the performance of CaMeRe on the clinical drug combination response datasets and the PDXs drug combination response datasets. We thus conclude that the effective synergistic integration of key components is crucial for enabling CaMeRe to make accurate predictions.

### E. Generalization results of CaMeRe in different domain scenarios

Adapting pretrained models to various scenarios is often an urgent necessity. To further evaluate the prediction performance of CaMeRe in different domain generalization scenarios, we compare CaMeRe with TCRP^+^ in following three distinct fine-tuning scenarios: (1) scenario 1, the dataset used to construct the support sets for meta-test tasks differs from that used for meta-training set; (2) scenario 2, there are discrepancies between the meta-training and meta-test support sets in terms of the number of cell lines, the number of samples in per cell line, or the number of unlabeled patients; (3) scenario 3, a new clinical/PDXs drug combination response dataset outside of the original meta-training datasets and meta-test dataset.

For the scenario 1, we retrieve cancer cell line drug synergy data from GDSC Combinations dataset ^4^, and choose the cancer cell lines corresponding to six tissues, including colon tissue (NCL=23), ovary tissue (NCL=16), breast tissue (NCL=19), lung tissue (NCL=58), brain tissue (NCL=24), and skin tissue (NCL=18). Then, we fine-tune the pretrained CaMeRe and TCRP^+^ using the new support sets (in which the cell lines samples are drawn from the GDSC Combinations dataset) to evaluate the performance of fine-tuned CaMeRe and fine-tuned TCRP^+^ on the test sets of meta-test tasks. The results of CaMeRe and TCRP^+^ in scenario 1 are shown in **Fig.4c**, from which we can see that the AUPR and BACC of CaMeRe are higher than TCRP^+^ on five tissue-specific clinical drug combination response datasets and four tissue-specific PDXs drug combination response datasets. These results indicate that the pretrained CaMeRe maintains better generalization ability, even when the support sets in meta-test tasks are built using a different dataset than the one used in the meta-training.

For the scenario 2, we leverage the dataset retrieved in scenario 1 to fine-tune the pretrained CaMeRe and TCRP^+^ with a series of distinct support set configurations. Here, the number (L) of cell lines takes the value from {2, 3, 4, 5, 6, 7, 8}, the number (K_C_) of samples in per cell line takes the value from {20, 40, 60, 80, 100, 120}, and the number (M) of unlabeled patients takes the value from {5, 6, 7, 8, 9, 10, 11, 12}. We then test the power of fine-tuned CaMeRe and TCRP^+^ with the test samples in meta-test sets, and the results of CaMeRe and TCRP^+^ in scenario 2 are shown in **Fig.4d**. From **Fig.4d**, we can see that in different support setting, the average AUROC of CaMeRe on five tissue-specific clinical drug combinations response datasets and four tissue-specific PDXs drug combinations response datasets are consistently higher than TCRP^+^. These results indicate that although there are discrepancies between the support set configurations in meta-training and meta-test task, the pretrained CaMeRe has superior generalization performance relative to TCRP^+^.

For the scenario 3, we first retrieve the clinical drug combination response data of bladder cancer patients (TP=39, TN=35) from TCGA ^30^, drug combination response data of pancreatic cancer PDXs (TP=26, TN=144) from previous paper ^33^, unlabeled clinical drug combination response data of bladder cancer patients (NUP=7) and pancreatic cancer patients (NUP=15) from TCGA, and drug synergy data of bladder cancer cell lines (NCL=8) and pancreatic cancer cell lines (NCL=13) from the GDSC Combinations dataset ^4^. Then, we establish a clinical drug combination response dataset with a focus on bladder tissue and a PDXs drug combination response dataset with a focus on pancreatic tissue, fine-tune and test the pretrained CaMeRe and TCRP^+^ on these two datasets, respectively. The average results of pretrained CaMeRe and pretrained TCRP^+^ in scenario 3 are shown in **Fig.4e**, from which we can see that AUROC and BACC of pretrained CaMeRe are higher than that of pretrained TCRP^+^ by 21.944% and 16.777% on bladder tissue dataset, 10.057% and 23.303% on pancreatic tissue dataset. These results indicate that despite facing new clinical/PDXs drug combination response dataset, the pretrained CaMeRe still exhibits stronger generalization ability compared to the pretrained TCRP+.

In short, above results across three scenarios demonstrate that our CaMeRe has stronger domain generalization than TCRP^+^.

### F. Interpretability of CaMeRe

Ensuring model interpretability is vital when applying prediction models to support medication decisions in personalized medicine. To assess the interpretability of CaMeRe in predicting specific drug combination response, we first used SHAP attribution decision method (see Methods for detail) to identify the top-rank features with the greatest contribution to the prediction results. Then, we systematically evaluated these features from four perspectives: (1) literature support; (2) prognostic relevance; (3) biological pathway enrichment; (4) consistency. Next, we take the combination of Paclitaxel plus Carboplatin from ovary tissue clinical drug combination response as a case study to demonstrate the interpretability of CaMeRe. The combination of Paclitaxel plus Carboplatin is a chemotherapy regimen (abbreviated as TC regimen) that is currently the most classic and standard first-line chemotherapy regimen for treating ovarian cancer ^34, 35^.

1. Literature support. **Fig.5a** shows the expression levels and the SHAP values of top 30 genes with greatest contributions in predicting the combination response of ovarian cancer patients to TC regimen. Among these top 30 genes, some of them have been reported to be closely associated with ovarian cancer or TC regimen in literatures. For example, FANCA (Fanconi Anemia Complementation Group A, Top 15 gene) is a core gene in the Fanconi anemia pathway ^36^, and some studies have shown that the low expression of FANCA is significantly associated with longer overall survival and greater sensitivity of ovarian cancer to platinum-based drug (such as Carboplatin) ^37^. RAD51C (RAD51 Paralog C, top 22 gene) is responsible for repairing double-strand DNA breaks and maintaining genomic stability ^38^, and it is recognized as a key biomarker for evaluating homologous recombination deficiency (HRD) status ^39^, and also used to predict sensitivity to platinum-based (such as carboplatin) chemotherapy ^40, 41^. BUB1B (BUB1 Mitotic Checkpoint Serine/Threonine Kinase B, top 26 gene) has been identified as a spindle assembly checkpoint gene associated with paclitaxel resistance ^42^ and high expression of BUB1B is significantly associated with poor prognosis in ovarian cancer, promoting tumor cell survival by facilitating mitotic progression and enhancing anti-apoptotic capacity ^43, 44^. DDIT4 (DNA Damage-Inducible Transcript 4, top 5 gene) is a negative regulator of the mTOR pathway, and it is closely associated with stress response, drug resistance, apoptosis, and tumor progression in ovarian cancer ^45, 46^. High expression of DDIT4 can activate autophagy and attenuate paclitaxel-induced apoptosis ^47^. MELK (Maternal Embryonic Leucine Zipper Kinase, top 28 gene) is a cancer stem cell maintenance factor in multiple tumors (including ovarian cancer, and cervical cancer), and its abnormal activation and overexpression are frequently associated with tumor progression, poor prognosis and chemotherapy resistance, particularly to paclitaxel ^48, 49^. Above analysis indicates that the genes with the greatest contributions identified by CaMeRe are closely correlated with the responses of ovarian cancer patients to TC treatment.
2. Prognostic relevance. To explore the relevance between gene expression levels and the survival period of patients treated with TC regimen, we collected the clinical data of ovary cancer patients receiving TC regimen treatment from TCGA database, and then stratified these patients (*i.e*., TCGA-OV cohort) into two groups based on the median expression levels of genes for performing Kaplan-Meier survival analysis. As shown in **Fig.5b** and **Fig.S10** (in Supplementary Information), we found that the expression levels of some genes in the top 150 genes are significantly associated with survival times in ovary cancer patients treated with TC regimen treatment. For example, at a survival rate of 0.5, the survival time of patients in DDIT4 low expression group (*i.e*., 37.6 months) is significantly (log-rank P-value < 0.0001) higher than that of patients in DDIT4 high expression group (*i.e*., 21 months); the survival time of patients in SFN low expression group (*i.e*., 36.7 months) is significantly (log-rank P-value=0.0044) higher than that of patients in SFN high expression group (*i.e*., 23 months); the survival time of patients in NPU93 low expression group (*i.e*., 38.6 months) is significantly (log-rank P-value=0.027) higher than that of patients in NPU93 high expression group (*i.e*., 21.3 months); the survival time of patients in DNAJ32 low expression group (*i.e*., 38.6 months) is significantly (log-rank P-value < 0.0001) higher than that of patients in DNAJ32 high expression group (*i.e*., 21.3 months). The prognostic survival analysis results indicate that CaMeRe can identify the genes whose expression levels are significantly associated with survival times in ovarian cancer patients receiving TC therapy.
3. Biological pathway enrichment. We also conducted enrichment analysis on the 150 top-rank genes identified by CaMeRe, including two kinds of gene enrichment analysis on Gene Ontology: Biological Process (GO: BP) and Kyoto Encyclopedia of Genes and Genomes (KEGG) pathway, as shown in **Figs.5c-5d**. From the hierarchical clustering results of the enriched GO: BP in **Fig.5c**, we found that the top-rank genes are significantly enriched in the biological processes that may affect the efficacy of TC regimen in ovarian cancer patients. For instance, Nuclear division cluster (consist of ‘Regulation of cell cycle’ term, ‘Mitotic cell cycle process’ term, ‘Sister chromatid segregation’, etc.) is directly related to the mechanism of paclitaxel directly interfering with microtubules and mitotic process ^50^; Regulation of molecular function cluster (consist of ‘Cellular response to stress’ term, ‘Apoptotic signaling pathway’ term, ‘Regulation of catalytic activity’ term, etc.) is directly related to the sensitivity of cells to recognize carboplatin induced DNA damage and apoptosis ^51^; Organic substance metabolic process cluster (consist of ‘Nitrogen compound metabolic process’ term, ‘Small molecule metabolic process’ term, ‘Phosphorus metabolic process’ term, etc.) reflects whether tumor cells have the ability to cope with drug pressure ^52^. Immune system development cluster (consist of ‘T cell homeostasis’ term, ‘Immune system development’ term, ‘Enzyme-linked receptor protein signaling’ term, etc.) plays abundant tumor-infiltrating lymphocytes and intact immune system function, and is associated with improved sensitivity to paclitaxel-carboplatin therapy and favorable prognosis ^53, 54^; Lymphocyte apoptotic process cluster exerts immunosuppression, which may further compromise treatment efficacy ^55^. These GOBP enrichment results suggest that our CaMeRe can successfully capture functionally relevant genes that mediate the effects of combination therapy. From the KEGG pathway enrichment results in **Fig.5d**, we found that the top-rank genes are significantly enriched in these pathways related to the efficacy of TC regimen in ovarian cancer patients. For example, Platinum drug resistance pathway is related with carboplatin resistance ^56^; Cell cycle pathway is the site of action of paclitaxel (G2/M blockade) ^57^; P53 signaling pathway coordinates DNA damage and apoptosis, responds to carboplatin ^58^; Apoptosis pathway is the endpoint pathway of the combination action of paclitaxel and carboplatin; PI3K-Akt signaling pathway regulates cell survival and resistance through anti-apoptotic signaling and metabolic adaptation, and is widely regarded as one of the core pathways for paclitaxel and platinum drug resistance ^59^. Above KEGG pathway enrichment results show that CaMeRe can identify pathways involved in the response to paclitaxel and carboplatin combination therapy in ovarian cancer patients, thereby enhancing its interpretability.
4. Consistency. We first used the trained CaMeRe to obtain three gene sets (each containing the top 150 genes) from three datasets: 10 ovarian cancer cell lines, 50 unlabeled ovarian patients, and 51 labeled ovarian patients, and then draw Venn diagram (**Fig.5e**) for these three gene sets. As shown in **Fig.5e**, 46 genes overlap across cell lines, unlabeled patients, and labeled patients; 107 genes overlap between unlabeled patients and labeled patients; 56 genes overlap between cancer cell lines and labeled patients; 56 genes overlap between cancer cell lines and unlabeled patients. We also analysis the correlation direction among three top-gene sets by drawing bubble plot (**Fig.5f**) and scatter plots (**Figs.5g-5j, Fig.S11**), where the correlation direction between gene expression value and its corresponding SHAP value is determined by the sign of Pearson correlation coefficient, and the contribution of each gene feature is measured by the mean absolute value of its corresponding SHAP value across all samples in specific dataset. From **Figs.5f-5j** and **Fig.S11**, we can see that many top-rank genes have same direction in cell lines, unlabeled patients, and patients. These results indicate that our CaMeRe can capture the consistency features among different data source domains, thereby enhancing the consistency and robustness of interpretation.

**Fig 5.**
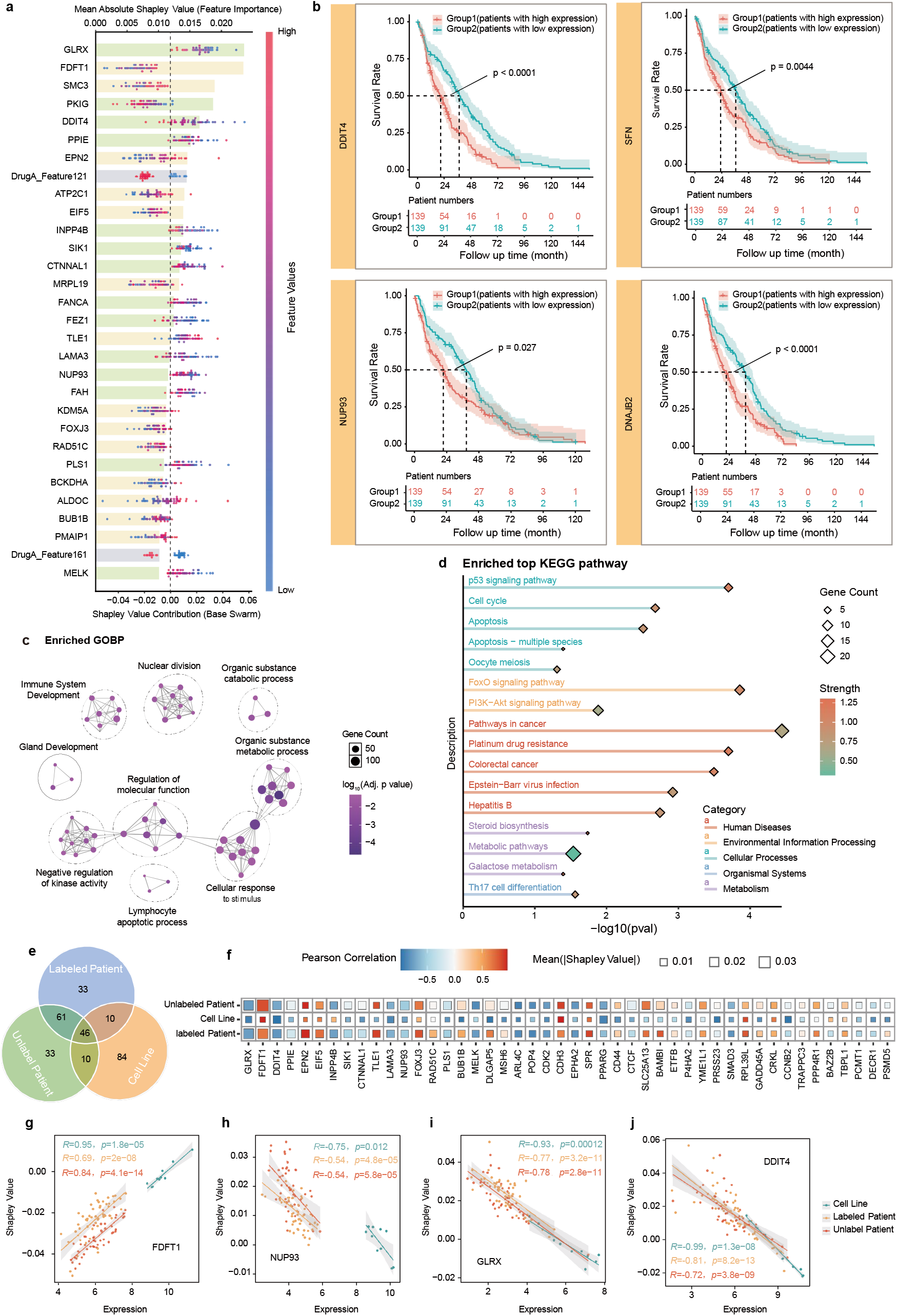
Interpretable results of CaMeRe in predicting the response of ovary cancer patients to TC regimen (*i.e*., Paclitaxel plus Carboplatin combination therapy). **a**, SHAP plots of 30 features with greatest contributions in predicting the combination response. Here, the bar plot shows the average contribution (*i.e*., mean absolute SHAP values) of each feature on the labeled samples from all ovarian cancer patients receiving TC regimen treatment; the bee swarm plot illustrates the distribution of the SHAP values, and each dot corresponds to one patient. **b**, Kaplan-Meier curves of Disease-Free Survival, in which the ovary cancer patients receiving TC regimen treatment are stratified into two groups based on the median expression levels of gene DDIT4, SFN, NUP93 and DNAJB2. **c**, The top GO: BP enrichment terms for the 150 genes with greatest contribution in predicting the response of ovary patients to TC regimen. Each node in the graph represents a pathway, where the color indicates the statistical significance (Benjamini-Hochberg correction hypergeometric test) of BP enrichment; each edge in the graph indicates the Jaccard similarity between two nodes. **d**, The top KEGG pathway enrichment terms for the 150 genes with greatest contribution. **e**, Venn diagram for illustrating the overlap of 150 genes with the greatest contribution in predicting the response of three different groups (*i.e*., 10 cancer cell lines, 50 unlabeled patients, and 51 labeled patient) to the TC regimen. **f**, Bubble plot of overlap genes with the greatest contribution in three groups. One bubble corresponds to a gene on X-axis. The size and color intensity of bubbles represent the average contribution and the Pearson correlation between Shapley values and expression levels of each gene, respectively. **g-j**, Scatter plots for visualizing the correlation between Shapley values and expression levels of gene FDFT1(**g**), NUP93(**h**), GLRX (**i**) and DDIT4 (**j**), respectively. One dot represents a cell line/unlabeled patient/labeled patient, and the dots are grouped into the three groups.

### G. The potential causal genes associated with drug combination response

To explore the potential causal genes associated with specific drug combination response, which is crucial for dissecting resistance pathways and guiding individualized therapy, we introduce the causal Shapley value insight ^60, 61^ into the backdoor adjustment block of CaMeRe to identify the top-ranked features that contribute most to the counterfactual response by intervening one gene expression level and forcing all other genes to remain unchanged in expression levels. These feature genes are called potential ‘causal’ genes (the inputs to the causal feature encoder) and ‘domain-specific’ genes (the inputs to the domain-specific feature encoder).

Considering that the combination of Encorafenib and Binimetinib (abbreviated as E+B regimen) is a targeted therapy regimen approved by FDA for the treatment of patients with unresectable or metastatic melanoma harboring BRAF V600E/V600K mutations ^62^, we take E+B regimen as a case study to identify the potential ‘causal’ genes. We first downloaded the PDXs E+B combination response dataset from skin tissue, in which 32 PDX models (*i.e*., 3 responses and 29 non-responses) received treatment with E+B regimen, and then constructed five counterfactual scenarios (*i.e*., Pdx_B1, Pdx_B2, Pdx_B3, Pdx_B4 and Pdx_B5) in CaMeRe backdoor adjustment block for identifying the potential ‘causal’ genes and ‘domain-specific’ genes via causal Shapley values, and also verified whether these potential ‘causal’ genes and ‘domain-specific’ genes meet the necessity conditions of stable causal factors and domain-specific response-modulating factors from three complementary perspectives ^63, 64^: (1) Do the potential ‘causal’ genes contribute more in predicting individual response to E+B regimen? (2) Are the contributions of potential ‘causal’ genes stable and the contribution of potential ‘domain-specific’ genes changing under different interventions? (3) Are the pathways enriched by potential ‘causal’ genes consistent with the drug combination target pathways? The analysis results of potential ‘causal’ genes and potential ‘domain-specific’ genes are shown in **Fig.S12**. From **Fig.S12**, we can see that i) the contributions (*i.e*., mean absolute Shapley value) of the potential ‘causal’ genes in predicting PDXs response to E+B regimen are significantly higher than the potential ‘domain-specific’ genes (**Fig.S12a**, ANOVA Test, *P* value < 2E-16); ii) the potential ‘causal’ genes have significant contribution consistency in five counterfactual scenarios (**Fig.S12b**, Chatterjee’s rank correlation Test, *P* value < 0.005), while potential ‘domain-specific’ genes exhibit considerable contribution heterogeneity in five counterfactual scenarios (**Fig.S12c**, Chatterjee’s rank correlation Test, P value > 0.05); iii) many KEGG pathways enriched by the potential ‘causal’ genes align with the important pathways targeted by Encorafenib and Binimetinib, including MAPK signaling pathway, PI3K-AKT signaling pathway, cell cycle, apoptosis, P53 signaling pathway(**Figs.S12d**-**e**), suggesting that our CaMeRe can successfully capture functionally consistent driving factors for therapeutic response. These results show that the potential ‘causal’/’domain-specific’ genes identified by CaMeRe meet the necessity conditions of stable causal factors and domain-specific response-modulating factors.

To further screen the genes that directly affect drug combination response from the potential ‘causal’ genes, we constructed a Directed Acyclic Graph (DAG) using gene regulatory network collected from TRRUST dataset ^65^ and OmniPath dataset ^66^, along with potential ‘causal’ genes and ‘domain-specific’ genes, and the drug combination responses variable **Y**. Based on the constructed DAG, we adopted Dagitty ^67^ to select the minimal adjustment set for each potential ‘causal’ gene, and then employed multiple classical causal estimation methods, including causal logistic regression model (CLRM) ^68^, DoubleML^69, 70^, cause forest ^71, 72^, instrumental variables regression (iverg) ^73, 74^, and regularized LASSO effects (rlassEffects) ^75, 76^, to estimate the causal effect of each potential ‘causal’ gene expression level on the response. As shown in **Fig.6**, for gene BCL2 (B-cell lymphoma 2), the causal effects of five causal estimation methods are all negative and statistically significant; for gene CCNA2 (Cyclin A2), the causal effects of five causal estimation methods are all negative, and the causal effects of three estimation methods are statistically significant; indicating that genes BCL2 and CCNA2 are significantly negatively related to PDXs response to E+B regimen. For gene TP53BP1 (Tumor Protein P53 Binding Protein 1), the causal effects of five causal estimation methods are all positive, and the causal effects of three estimation methods are statistically significant, indicating that gene TP53BP are significantly positively related to PDXs response to E+B regimen. These results suggest that BCL2, CCNA2, and TP53BP1 may be the candidate ‘causal’ genes in the response of PDX models to the E+B regimen.

**Fig 6.**
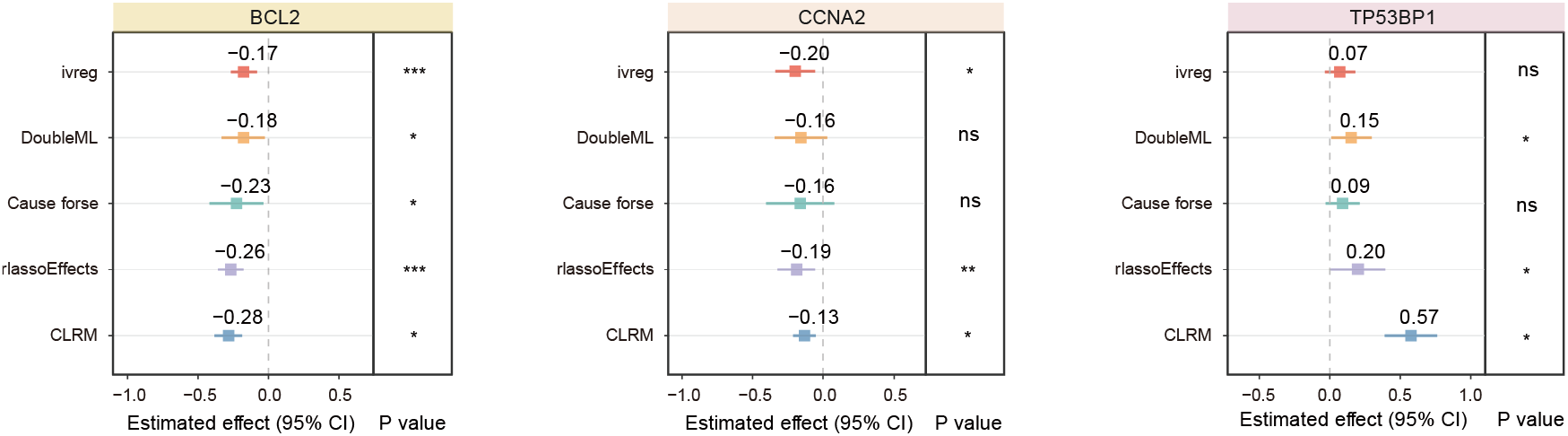
Forest plots for illustrating the estimated causal effects of ‘causal’ genes (*i.e*., BCL2, CCNA2, TP53BP1) on the response of skin cancer PDX models to the combination of Encorafenib and Binimetinib. The magnitude of the causal effect represents the amount of change in the response for a one-unit increase in one ‘causal’ gene expression, whereas expressions of minimal adjustment gene set for the ‘causal’ gene are fixed. The sign (+, −) of causal effect denotes whether there is a positive (+) or negative (−) correlation between ‘causal’ gene and response.

### H. Efficacy evaluation of CaMeRe in predicting large-scale unlabeled clinical patients to drug combination responses

To verify the effectiveness of CaMeRe in predicting large-scale unlabeled clinical patients to drug combination responses, we retrieved 26 drug combinations from the National Cancer Institute (https://www.cancer.gov/about-cancer/treatment/drugs, it compiles drug combinations approved by FDA for cancer therapy) and DrugComDB dataset ^77^ (it compiles potential drug combinations that have been extensively investigated in cancer therapy) and downloaded the gene expression profiles of 3,423 clinical patients from TCGA database for pharmacogenomic analysis, aiming to uncover: (1) whether the predicted drug combination responses are significantly associated with the expression of target genes or known biomarkers; (2) whether the high response samples are enriched into drug combination relevant pathways. The heatmap of the response scores (predicted by CaMeRe) of 3,423 patients to the 26 drug combinations is shown in **Fig.S13**, and the pharmacogenomic results of 26 drug combinations are presented in **Table S10**. From **Table S10**, we found that for 23 out of 26 drug combinations, the predicted drug combination response scores are significantly associated with the expression levels of their target genes or known biomarkers. Moreover, for 23 out of 26 drug combinations, high-response samples are significantly enriched in the pathways related to drug combination responses. For instance, for the combination of Cisplatin plus Gemcitabine, a standard chemotherapy regimen that has been extensively utilized in clinical practice across various solid tumors (particularly among patients with advanced-stage or surgically unresectable non-small cell lung cancer), the predicted drug combination response of lung cancer patients is significantly correlated with the expression of biomarker genes (**Fig.7a**), such as BRCA1/2, RRM1/2, CASP3, while these genes play crucial roles in lung cancer cell proliferation and nucleotide metabolism (*e.g*., RRM1/2), DNA repair mechanisms (*e.g*., BRCA1/2), and apoptosis (*e.g*., CASP3). More importantly, their expression levels are closely associated with the combination therapy efficacy of Cisplatin plus Gemcitabine for lung cancer ^78-80^.

**Fig 7.**
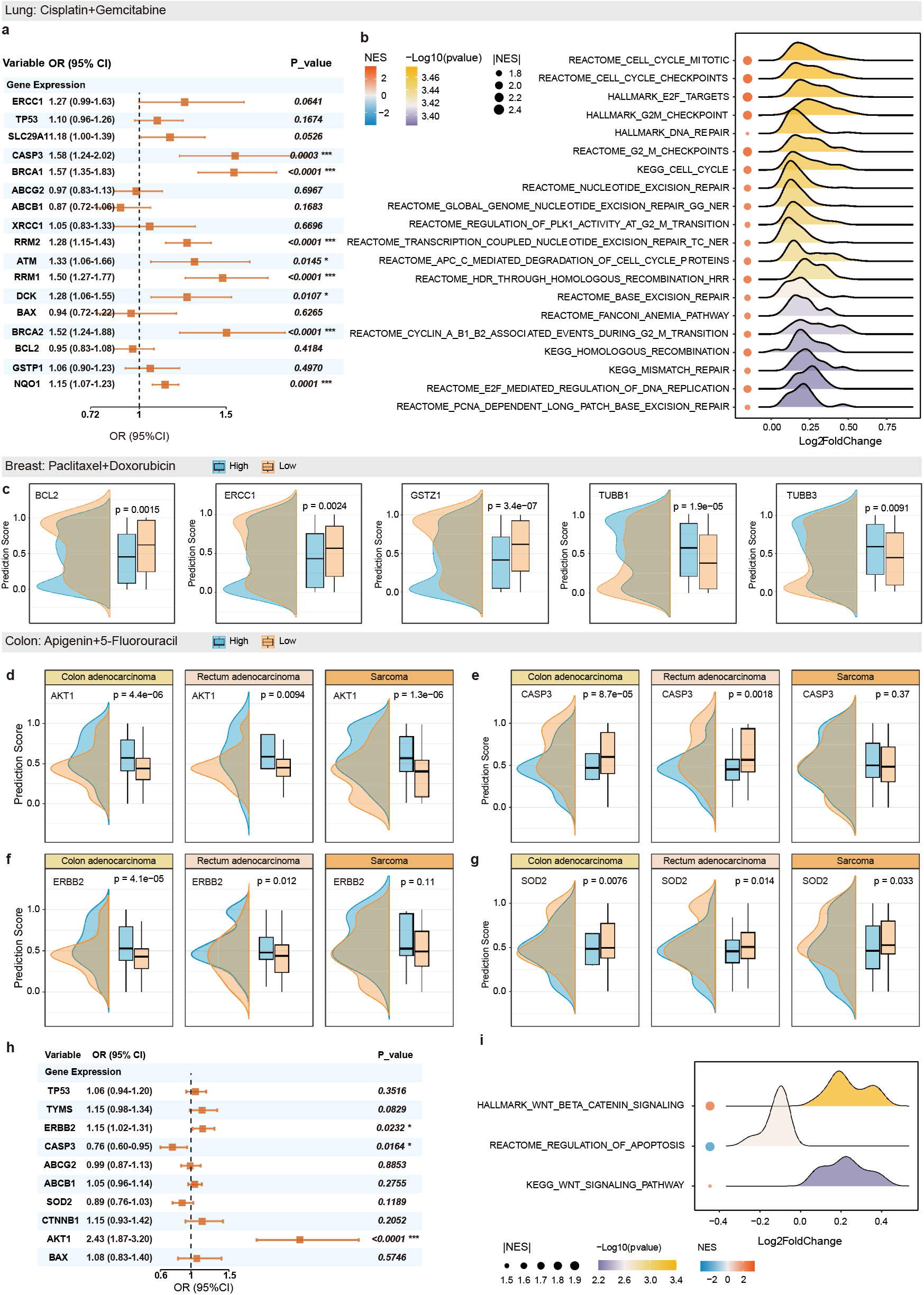
The pharmacogenomic results of CaMeRe on three drug combinations approved by FDA. **a**, Forest plot for showing the results of logistic regression of the predicted combination response scores (binarization based on median value) and known genes related to combination response of Cisplatin plus Gemcitabine for lung cancer patients. Each square represents the Odds Ratios (OR), and horizontal lines indicate the 95% confidence intervals (CIs). **b**, Bubble plot and Ridge plot for showing the known pathways (identified by GSEA) related to combination response, where the lung cancer patients are stratified based on the median predicted response score to the combination of Cisplatin plus Gemcitabine. The ridge illustrates the distribution of the differential expression levels (log_2_ fold change, log_2_FC) of genes within each pathway entry between the high-response and low-response groups, the color of ridges indicates the negative log-transformed significance (−log_10_P-value) of pathway enrichment. The color of the bubbles represents the normalized enrichment score (NES) of the corresponding pathway, a positive NES value represents pathway upregulated in the high response group, whereas a negative value represents pathway downregulated in the high response group, and the size of the bubbles denotes the absolute value of the normalized enrichment score (|NES|) of the corresponding pathway. **c**, Half-violin plots for illustrating the distribution of the predicted response scores to the combination of Doxorubicin plus Paclitaxel in breast cancer patients with high (above the median) or low (below the median) expression of one gene (*i.e*., BCL2, ERCC1, GSTZ1, TUBB1, and TUBB3) related to drug combination response. The BH-correction P-values are assessed by ANOVA test. **d-i**, The pharmacogenomic results of CaMeRe on 5-Fluorouracil and Apigenin combination for colon cancer patients.

In addition, according to the median predicted response scores of patients to the combination of Cisplatin plus Gemcitabine, we classify the lung cancer patients into a high response group and a low response group, then use GSEA to identify the pathways related to combination response (**Fig.7b**). From **Fig.7b**, we can see that the pathways of cell cycle mitotic (Reactome), E2F targets (Hallmark), homologous recombination (KEGG), and E2F mediated regulation of DNA replication (Reactome) are significantly enriched in high response groups, while these pathways are closely related to the pathogenesis of lung cancer. Specifically, these pathways are closely associated with the proliferation and repair mechanisms of lung tumor cells, and they also bear a strong relationship to lung cancer patients’ response to DNA-damaging treatment regiments, including the combination of Cisplatin plus Gemcitabine ^78, 81^.

For the combination of Doxorubicin plus Paclitaxel, a combination chemotherapy scheme that has been widely studied in solid tumors such as breast cancer, ovarian cancer, endometrial cancer, lung cancer, *etc*., the predicted drug combination response of breast cancer patient is significantly correlated with the expression of biomarker genes (**Fig.7c**), including BCL2, ERCC1, GSTZ1,TUBB1 and TUBB3; the predicted drug combination response of ovarian cancer patients is significantly associated with the expression of biomarker genes (**Fig.S14**), including CDK1, CCNB1, TUBB, ATM, GSTZ1, ERCC1, TOP2A. In various solid tumors such as breast cancer, ovarian cancer, and lung cancer, the expression levels of these genes correlate with target abundance/ tumor proliferation (*e.g*., TOP2A, TUBB, TUBB1/3), cell cycle activity (*e.g*., CDK1, CCNB1), and anti-apoptotic and DNA damage repair capabilities (*e.g*., BCL2, ERCC1, ATM, GSTZ1). These genes also influence response of cancer patients to the combined therapy of Doxorubicin plus Paclitaxel ^82-87^.

For the combination of 5-Fluorouracil and Apigenin, a potential combination chemotherapy scheme for colon cancer, the predicted drug combination response of colon cancer patients is significantly associated with the expression of biomarker genes (**Figs.7d-h**), such as AKT1, CASP3, ERBB2, SOD2, and some pathways are significantly enriched in high/low response group (**Fig.7i**). In colon cancer, the Wnt/β-catenin signaling pathway serves as a core driver pathway that not only promotes tumorigenesis but also enhances cancer cell proliferation and survival ^88^. Concurrently, the genes AKT1, ERBB2, CASP3, and SOD2 are also closely associated with tumor proliferation, progression, apoptosis, and oxidative stress responses. Moreover, these genes and pathways are closely related to the synergistic mechanism of Apigenin enhances the cytotoxic effects of 5-Fluorouracil ^89-91^.

The above results support the effectiveness of CaMeRe in predicting responses of large-scale unlabeled clinical cancer patients to drug combinations from the perspective of biological plausibility.

### I. Screening potential drug combinations for specific cancer with CaMeRe

To screen the potential drug combinations for specific cancer from large-scale patients and drug combinations, we downloaded the gene expression profiles of 3,423 cancer patients across 11 cancer cohorts (*i.e*., LUAD, LUSC, UCEC, OV, UCS, BRAC, COAD, SARC, READ, LGG, GBM) from TCGA ^30^, as well as the drug structure data of 95 chemotherapy drugs and 945 targeted drugs that constitute 542,080 drug combinations from PubChem dataset ^92^. Then, we used CaMeRe to predict the response scores of 3,423 patients to 542,080 drug combinations. The 3,423 patients to 542,080 drug combinations can form above 1.8 billion drug combination-patient pairs, as shown in **Fig.8a**. Based on the predicted response scores of cancer patients to drug combinations, we screened the potential drug combinations that are significantly enriched in high-response group of each cancer cohort. Specifically, we selected the drug combination-patient pairs with response scores above 0.9 for each cancer cohort, and employed hypergeometric test for enrichment analysis. The statistical results of potential drug combinations enriched in each cancer cohort are provided in Supplementary Information **Note 2** and **Fig. S15**.

**Fig 8.**
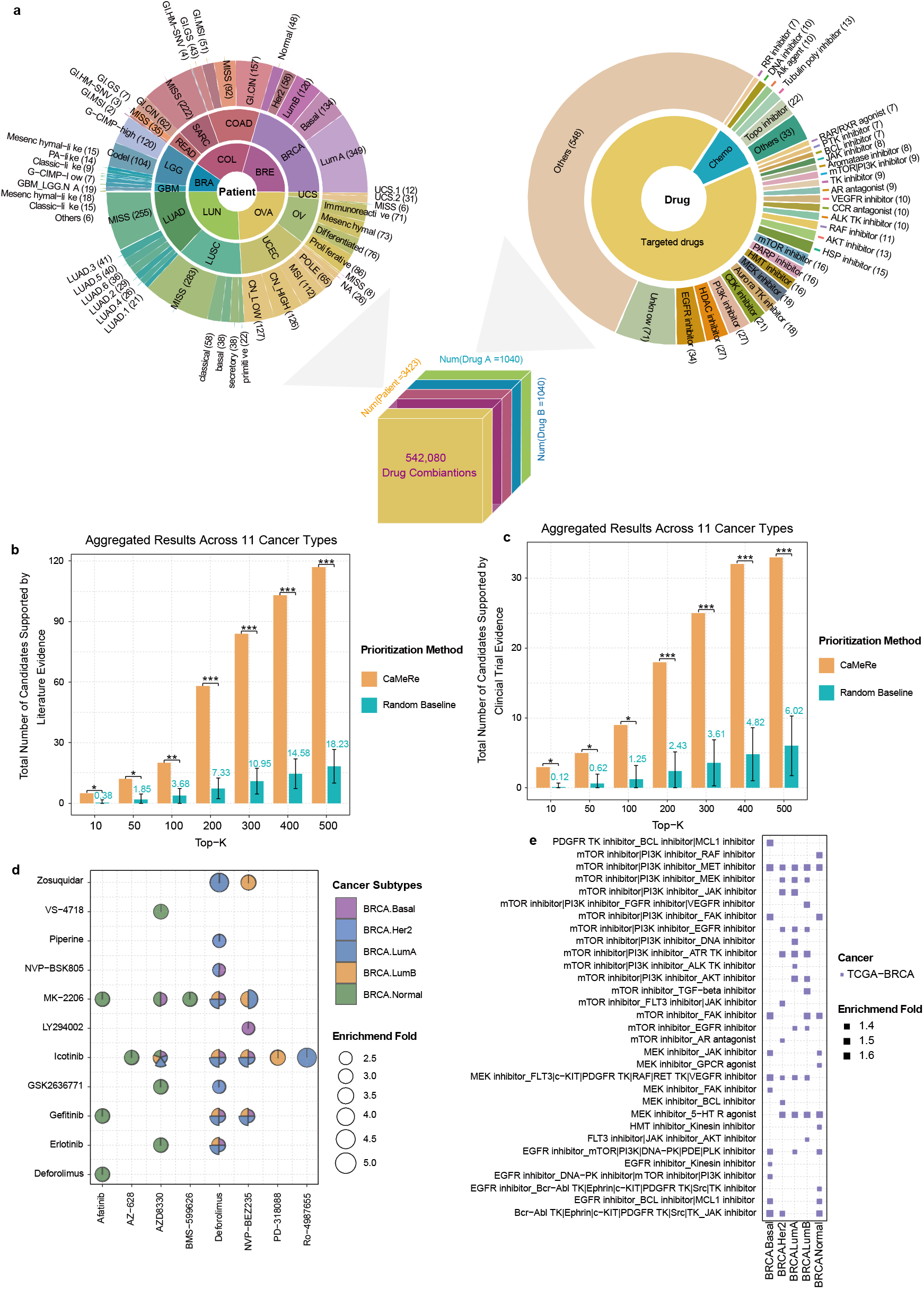
Screening results of CaMeRe on large-scale drug combinations for specific cancers. **a**, overview map of the cancer patients and drug combinations. The number in parentheses is the number of patients in each cancer subtype or the number of drugs annotated with the same mechanism of action. **b**-**c**, Bar plots showing the aggregated comparison results of evidence-support candidate counts 11 cancer types at different Top-*K* thresholds. **b**. Total number of literature evidence-supported candidates prioritized by CaMeRe and the random baseline across 11 cancer types. **c**. Total number of clinical trial evidence-supported candidates prioritized by CaMeRe and the random baseline across 11 cancer types. Random baseline bars represent the mean candidate counts, with error bars indicating the standard deviation, and numerical labels above the bars indicating the corresponding mean counts. Statistical significance is assessed using empirical permutation test with Benjamini-Hochberg correction, and significance levels are marked by asterisks. ‘ns’, not significant; ‘*’, BH-corrected P<0.05; ‘**’, BH-corrected P<0.01; ‘***’, BH-corrected P<0.001. **d**, Top 10 drug combinations significantly enriched in high-response (*i.e*., the predicted response scores above 0.9) groups of each breast cancer subtype. **e**, Top 10 MOA drug pairs significantly enriched in high-response groups (*i.e*., the predicted response scores above 0.9) of each breast cancer subtype. For **d**-**e**, enrichment significance is evaluated using hypergeometric tests, followed by Benjamini-Hochberg correction for multiple testing.

To systematically evaluate the capability of CaMeRe for large-scale drug combination screening, we first ranked the significantly enriched drug combinations using the enrichment fold and the percentage of high-response patients in each cancer cohort, and then performed a uniform literature search and clinical trials search for the top-*K* candidates (10, 50, 100, 200, 300, 400, 500) in each cancer cohort. For each top-*K* set, we quantified the number of candidates supported by published literature and registered clinical trials, respectively, and compared them with those of randomly selected *K* candidates. Random sampling was repeated 10,000 times to construct empirical baselines for assessing the statistical significance. The corresponding results are presented in **Fig.8b-c** and **Fig.S16**. As shown in **Fig.8b** and **Fig.8c**, for the aggregated analysis across cancer types, the top-*K* candidates prioritized by CaMeRe exhibit significantly greater literature support and clinical-trial support than the random baseline, with all empirical permutation P values below 0.05 after BH correction. Here, the aggregated random baseline was constructed by randomly sampling the same number of candidates within each cancer type 10,000 times and then aggregating the support counts across cancer types. Furthermore, as shown in **Fig. S16**, for the cancer-type-specific analysis, the top-*K* candidates prioritized by CaMeRe show significantly greater literature support in 9 cancer types and significantly greater clinical-trial support in 7 cancer types. In each of these cancer types, significance was observed at more than two top-*K* thresholds (empirical permutation P values below 0.05 after BH correction). Above these results indicate that CaMeRe can effectively prioritize drug-combination candidates with stronger existing evidence support for each cancer cohort from a large candidate space, highlighting its potential utility in large-scale drug-combination screening and prioritization.

Furthermore, for each cancer subtype cohort, we rank the significantly enriched drug combinations using the enrichment fold and the percentage of high-response patients in this subtype cohort. Top 10 drug combinations of BRCA cancer subtypes and other cancer (i.e., Ovary, Colon, Brain, Lung) subtypes are shown in **Fig.8d** and **Fig. S17**. From **Fig.8d** and **Fig. S17**, we can see that some drug combinations are enriched in more cancer subtypes, while some drug combinations are only enriched in a specific cancer subtype. For example, the combination of MK-2206 (AKT inhibitor) and Deforolimus (rapamycin-based mTOR1 inhibitor, also known as Ridaforolimus) is enriched in four BRCA subtypes (*i.e*., BRCA.LumA, BRCA.LumB, BRCA.Her2, BRCA.Basal). The phase I clinical study (NCT01295632) demonstrates that the combination of MK-2206 and Deforolimus exhibits biological activity in subsets of patients with advanced breast cancer ^93^. The combination of Gefitinib/Erlotinib (EGFR inhibitor) and Deforolimus is enriched in six non-small cell lung cancer subtypes. Some literature studies show that the dual inhibition of the EGRF/mTOR pathways may be a promising approach to treat EFGR wild-type NSCLC ^94^. Moreover, there are two clinical trials (NCT00456833, NCT00096486) assessing the efficacy and safety of rapamycin-based mTOR inhibitor Everolimus combined with Erlotinib or Gefitinib in patients with non-small cell lung cancer (NSCLC), respectively, finding that the combination of Everolimus and Erlotinib performs better in disease control rate without new safety concerns, compared with Erlotinib alone ^95^, and 13% patients have response to the combination of Everolimus and Gefitinib ^96^. These results suggest that CaMeRe can prioritize clinically supported/relevant candidate drug combinations for each cancer type, while capturing heterogeneity in candidate prioritization across different subtypes within the same cancer type. In addition, we investigated whether the enriched drug combinations are related to clinical features (*i.e*., tumor stage and patient age) by calculating the proportion of high response patients in each tumor stage cohort and age cohort. The results of top 5 enriched drug combinations for the three subtypes of BRCA cancer are show in **Fig. S18**, from which we can see that for one enriched drug combination, the proportion of high response patients is related to the tumor stage and age, that is, the proportion of high response patients is relatively high in some tumor (or age) stages and relatively low in other stages. For example, the combination of Deforolimus and Gefitinib has higher proportion of patients with high response scores at early to intermediate stage (67%) and young age (76%) for BRCA.Her2 subtype.

In view of that drug mechanism of action (MoA) target annotations can provide mechanistic understanding of drug action at molecular level, and relate protein targets to human disease and symptoms, we group drugs into different MoA clusters (*i.e*., mTOR|PI3K inhibitor, AKT inhibitor, AR antagonist, 5-HT R agonist, etc.) based on the annotation information of drug MoA targets, and then map the significantly enriched drug combinations to their corresponding MoA clusters to form MoA drug pairs, for analyzing which MoA drug pairs are significantly enriched response in specific cancer subtypes. The metrics of enrichment significance, fold enrichment, and the proportion of high response patients are used to rank the MoA drug pairs significantly enriched in each cancer subtype. The results of top 10 MoA drug pairs significantly enriched in each cancer subtype are shown in **Fig.8e** and **Fig.S19**, from which we can see that some MoA drug pairs are enriched in more cancer subtypes, while some MoA drug pairs are only enriched in a specific cancer subtype. For example, the MoA drug pair of mTOR|PI3K inhibitor (*e.g*., NVP-BEZ235, GSK2126458, PF-04691502, Voxtalisib) and MEK inhibitor (*e.g*., MEK162, PD-0325901, Selumetinib, Trametinib, AS-703026) is enriched in three subtypes of BRCA (*i.e*., BRCA.Her2, BRCA.LumA, BRCA.LumB), and some studies have shown that in breast cancer, the PI3K-AKT-mTOR and RAS-RAF-MEK-ERK pathways are two dominant tumor growth-promoting pathways, and the combination of mTOR|PI3K inhibitor and MEK inhibitor simultaneously blocks the PI3K-AKT-mTOR and RAF-MEK-ERK pathways and their mutual feedback loops, thereby producing a potent synergistic antitumor effect in breast cancer ^97, 98^. Several phase I clinical trials also investigated the combination of mTOR|PI3K inhibitor and MEK inhibitor for the treatment of breast cancer, such as, NVP-BEZ235 plus MEK162 (NCT01337765), Voxtalisib plus AS-703026 (NCT01390818) ^97^. The combination of MEK inhibitor and multi-kinase inhibitor sorafenib is enriched in four subtypes of BRCA, including BRCA.Basal (same as Triple-Negative Breast Cancer). In triple-negative breast cancer models, the combination of MEK inhibitor (*e.g*., AZD6244, also known as Selumetinib) and sorafenib exerts synergistic antitumor effects by concurrently targeting multiple kinases, thereby enhancing apoptosis and promoting more pronounced tumor regression than either agent alone ^99, 100^. The combination of EGFR inhibitor and BCL|MCL1 inhibitor is enriched in 5 subtypes of lung cancer (*i.e*., LUAD.3, LUAD.4, LUAD.5, LUAD.6, LUSC.classical). Dual inhibition of EGFR and anti-apoptotic BCL-2 family proteins (including MCL-1) can effectively overcome resistance to EGFR-targeted therapy in non-small cell lung cancer ^101^. Compared with EGFR inhibitor monotherapy, the combination of EGFR inhibitors with BCL2|MCL1 inhibitor (*e.g*., AT-101, also known as Gossypol) significantly delays lung tumor growth and without producing notable toxic effects ^101-103^. Moreover, in a phase II clinical trial (NCT00988169) for assessing the combination of EGFR inhibitor Erlotinib and BCL2|MCL1 inhibitor AT-101 in treatment-naive patients with EGFR-mutation advanced NSCLC, among six enrolled patients, one achieved a partial response, one had a minor response, and three exhibited stable disease ^104, 105^. The combination of ALK TK inhibitor and HSP inhibitor is enriched in 2 subtypes of lung cancer (*i.e*., LUAD.5, LUSC.classical). In ALK-rearranged NSCLC, the combination of ALK TK inhibitor and HSP inhibitor has emerged as a key synergistic strategy to overcome both primary and acquired resistance to ALK tyrosine kinase inhibitors ^106, 107^. Moreover, in a phase I clinical trial (NCT01579994), the efficacy of the HSP inhibitor ganetespib combined with the ALK-TK inhibitor crizotinib was evaluated in twelve patients with ALK-rearranged metastatic NSCLC, and it was found that 67% of patients achieved a partial response, confirming the feasibility and potential clinical benefit of this combination therapy (*i.e*., ALK TK inhibitor plus HSP inhibitor) ^108^. These results suggesting that CaMeRe can prioritize clinically relevant MoA drug pairs for each cancer type and capture subtype-dependent heterogeneity in enriched MoA patterns.

## Discussion

Despite advances in precision oncology, large-scale predicting combination therapeutic response in cancer patients remains challenging, due to tumor heterogeneity, the lack of robust biomarkers, and small clinical combination therapeutic response data available. To address these challenges, we propose a novel CaMeRe framework in this work to predict clinical drug-combination response. The experimental results demonstrate that our CaMeRe not only exhibits superior predictive performance compared to several representative methods, but also has good interpretability and the capability to screen drug combinations for specific clinical patients on a large scale.

The main difference between our CaMeRe and other methods is that it can train a two-level domain generalization model to generalize across inter-tissue domain and intra-tissue domain by using a small number of labeled clinical drug-combination response samples, with the assistance of labeled preclinical (*i.e*., cell lines) drug synergy samples, and a small number of unlabeled clinical drug-combination response samples. Specifically, (1) When stable causal factors and domain-specific response-modulating factors are invisible, CaMeRe designs a domain-invariant causal representation learning module based on the invariant information bottleneck theory and causal intervention invariance principle, to extract drug combination specific causal latent representations from raw inputs. (2) Since explicit discrete domain labels are unavailable, CaMeRe recast the supervised multi-domain discriminator as an unsupervised multi-domain discriminator driven by Wasserstein distance. Through joint adversarial training with the drug combination response block and the backdoor adjustment block, the feature embedding module can learn causal latent representations and domain-specific latent representations that are identical to the properties of stable causal factors and domain-specific response-modulating factors, respectively. (3) Due to limited sample sizes of cell lines, unlabeled patients, and patients corresponding to each drug combination, CaMeRe builds a meta-learning framework to optimize the parameters in the domain-invariant causal representation learning model. In the inner-loop learning process of CaMeRe, the base-learner first simulates a multi-domain generalization scenario corresponding to a specific drug combination, and then optimizes the parameters of base-learner for generalizing to each cell line or unlabeled patient in a specific tissue. In the outer-loop learning process of CaMeRe, the performance of base-learner is tested on the labeled patients, and the mean loss in multiple tissues is used to optimize the parameters of meta-learner for generalizing to multiple tissues. In addition, taking the cell lines different from the support set as auxiliary domains for the query set can reduce overfitting in the out-loop learning process, which is caused by the small size of the labeled clinical drug combination response data in each tissue. (4) Because the dataset exhibits an imbalanced label distribution in cell lines and patients, the Balanced Softmax function is utilized to correct the category prior bias of Softmax from a probabilistic perspective.

Under the constraints induced by our structural causal model, CaMeRe is encouraged to learn the mechanisms that remain invariant across individuals (*e.g*., cell lines or patients) and genuinely influence drug combination response, thereby endowing the CaMeRe with good interpretability. Based on the results of permutation SHAP analysis, we found that CaMeRe can capture important genes associated with the clinical response of ovarian cancer patients to drug combination of paclitaxel and carboplatin. Notably, DDIT4 gene, one of the major contributors to the prediction performance, has previously been reported in the literature to be associated with clinical response to combination therapy. Our survival analysis also shows that DDIT4 expression is significantly associated with survival time of ovarian cancer patients receiving combination therapy. Moreover, DDIT4 consistently ranks among the top contributors across diverse domains (cell lines, unlabeled patients, labeled patients), and its expression levels show a negative correlation with the prediction results in all domains, indicating that DDIT4 may be an important drug-resistant gene.

Different from other existing domain generalization methods for drug response prediction (*e.g*., Verlome), our CaMeRe has a backdoor-adjustment block which is designed to reduce confounding effects induced by ‘domain-specific’ genes), and it may obscure the underlying causal relationships between ‘causal’ genes and drug combination responses. Basing on the causal SHAP analysis results, potential ‘causal’ genes show stronger and more stable contributions than potential ‘domain-specific’ genes in predicting PDX response to the combination of Encorafenib and Binimetinib across different interventions. In addition, the pathways enriched by these potential ‘causal’ genes are consistent with the known target pathways of the drug combination. Above these findings support the biological plausibility of the ‘causal’/’domain-specific’ distinction made by CaMeRe, suggesting that CaMeRe can identify genes with potential causal relevance while mitigate the influence of domain-specific response-modulating factors in response prediction. Importantly, with the help of gene regulatory network and classical causal estimation methods, our findings reveal that the expression levels of genes BCL2, CCNA2 exhibit significant negative causal effects on the drug combination response, whereas the expression level of gene TP53BP1 shows significant positive causal effects. These direction-specific causal effects further support the biological interpretability of CaMeRe, suggesting that CaMeRe can identify candidate genes potentially linked to resistance or sensitivity in combination therapy. Nevertheless, these genes should still be regarded as putative causal factors, and their mechanistic roles remain to be validated experimentally.

In addition, unlike some existing clinical drug combination recommendation methods, CaMeRe takes the gene expression profiles of the patients and the molecular structures of the drug combinations as inputs, enabling it to predict response scores of clinical patients to any drug combinations with known molecular structure on a large scale. For example, for an 85-year-old male patient (ID: TCGA-A6-2671) with Stage IV Chromosomal Instability Colon Adenocarcinoma, CaMeRe outputs 0.4406 response score to the combination therapy of 5-fluorouracil and Leucovorin, and the result is consistent with the TCGA record, which indicate that the patient exhibited Clinical Progressive Disease following treatment with this drug combination. For a 62-year-old male patient (ID: TCGA-AD-6965) with stage III Chromosomal Instability Colon Adenocarcinoma, CaMeRe outputs 0.9609 response score to the combination therapy of 5-Fluorouracil and Leucovorin, which is in agreement with the TCGA clinical data, where this patient was also documented to have Complete Response following treatment with the drug combination. Pharmacogenomic analysis shows that the clinical drug combination response scores predicted by CaMeRe are significantly correlated with response biomarkers of drug combinations, as well as with the pathways associated with the drug combinations.

Based on the response scores predicted by CaMeRe for over 1.8 billion drug combination-patient pairs, we screened a broad space of drug combinations (including 542,080 drug combinations) to identify the drug combinations significantly enriched in 45 cancer subtypes, such as the combination of MK-2206 and Deforolimus enriched in BRCA.LumA subtype, and we also observed that the combination of PI3K|mTOR inhibitor (*e.g*., NVP-BEZ235, PF-05212384) and AKT inhibitor (*e.g*., MK-2206, AZD5363, GDC-0068) is enriched in BRCA.LumA/LumB subtypes. Mechanistically, these significantly enriched drug combinations predominantly target the PI3K-AKT-mTOR pathway, which is closely related to LumA/LumB breast cancer. Moreover, some studies ^109, 110^ have demonstrated that single-agent targeting of PI3K or mTOR often leads to compensatory AKT reactivation, whereas dual inhibition effectively suppresses downstream signaling and induces cell apoptosis in LumA/LumB breast cancer.

Although our CaMeRe can effectively predict drug combination response of clinical patients on a large scale and capture the ‘causal’ genes associated with the drug combination response, it also has the following limitations. First, due to the limited available data on clinical drug combination response, CaMeRe stratifies the cell lines, patients, PDXs according to their primary tumor tissue, and encourages the model to capture causal mechanisms of drug-combination response that are stable and transferable within each tissue type. CaMeRe has narrowed the scope of domain invariance compared to other drug response methods that constrain consistent mechanisms across all tissues, but further refinement of constraints (*e.g*., constraining consistent response mechanisms for drug combinations within specific cancer type) can improve predictive performance and uncover more granular causal mechanisms. Second, to mitigate the potential overfitting associated with high-dimensionality and small sample size, CaMeRe only uses the transcriptome data to characterize individuals. However, when using pharmacogenomic methods to evaluate the efficacy of CaMeRe in predicting large-scale unlabeled clinical patients to drug combination responses, the results show that the predicted drug combination response scores are significantly correlated with the expression levels of marker genes, but not with the mutational status of the marker genes. In the further work, we should consider introducing multi-omics data to effectively predict drug combination response and comprehensively explain the causal mechanisms of individual differences in drug combination responses.

Overall, CaMeRe demonstrates that integrating causal reasoning with meta-representation learning can significantly improve the prediction performance of drug combination response, highlighting how machine intelligence approaches effectively bridge preclinical data with personalized clinical decision-making, thereby promoting the rapid development of precision oncology.

## Methods

### I. Notations and Problem Formulation

Let 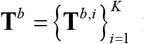 represent the set of base tissues, 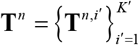 represent the set of new tissues, and these two sets do not intersect, *i.e*., **T**^*b*^ ⋂ **T**^*n*^ =Ø. Each base tissue **T**^*b,I*^ contains cell line samples derived from domains 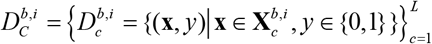, few unlabeled patient samples derived from domains 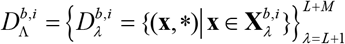, and few labeled patient samples derived from domains 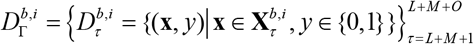. Where 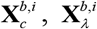, and 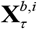 denote the feature vectors of cell lines, unlabeled patients, and labeled patients in the *i*-th base tissue, respectively; *y* denotes the drug combination response of cell lines/patients. In particular, the domain 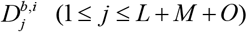 is drawn from the sample distribution 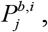, and 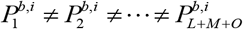. Analogously, 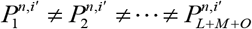, and 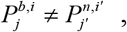, (1 ≤ *j*′ ≤ *L*′ + *M* ′ + *O*′, 1 ≤ *i* ≤ *K*, 1 ≤ *i*′ ≤ *K*′).

We model the patient’s drug combination response prediction in new tissue **T**^*n,i*′^ as a small sample bi-level domain generalization (DG) task. Generally, DG setting assumes the existence of domain-invariant patterns *S*^*n,i*′^ in inputs 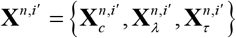, which can be extracted to learn a label predictor *f* (**θ**^*n,i*′^) that performs well across the seen domains and unseen domains. Unlike domain adaptation, DG assumes that the target domain data or explicit knowledge about the target distribution 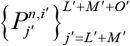 are unavailable during training. Therefore, given a set of base tissues **T**^*b*^ and their samples drawn from domains 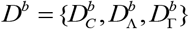, and a new tissue **T**^*n,i*^′ and its samples drawn from domains 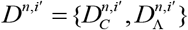, the goal is to quickly learn a function *f* (**θ**^*n,i*^) : **X**^*n,i*′^ → {0,1} with the samples drawn from 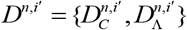, after learning a meta-function **F** (**θ**) : **X** →{0,1} with the samples drawn from *D*^*b*^, for predicting drug combination response of clinical patients in new tissue **T**^*n,i*′^.

### II. Structural Causal Model for Drug Combination Response

We formulate the transfer of drug-combination response prediction from cell lines to patient samples as a cross-system mechanism transfer problem, and employ a structural causal model to formalize the sample-level data-generating process. Specifically, let **E** = {cell line_1_,⃛, cell line_L_, patient_1_,, patient_(M+O)_} denote the domain indicator. Let **X** denote the observed gene expression profile, and **Y** denote the observed drug-combination response outcome, with **Y** =1 indicating response and **Y** = 0 indicating non-response. Let **D** denote the treatment-related variables, here it is represented by the joint molecular graph representation of drug combination. Let **S** denote the latent tumor-intrinsic biological states that are directly relevant to drug-combination response, such as pathway dependency, target activation, DNA repair deficiency or apoptotic priming. Let **V** denote the host-context variables and latent measurement-related variables, such as microenvironmental context, stromal composition, tissue architecture, systemic physiological state, platform effects, batch effect, preprocessing variation and tumor purity. Then, we model the data generation process via the following structural equations:

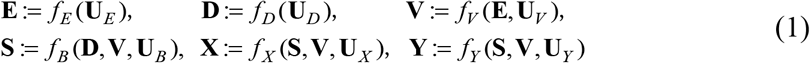

where **U**= {**U**_E_, **U**_D_, **U**_V_, **U**_B_, **U**_X_, **U**_Y_} are the corresponding exogenous variables. To make the structural dependencies more explicit, we rewrite above system of structural equations as a directed acyclic graph (DAG), as illustrated in **Fig.1a**. Specifically, whenever a variable appears on the right-hand side of the structural equation of another variable, a directed edge is induced from the former to the latter in the corresponding causal graph. The corresponding directed relations are as follows:

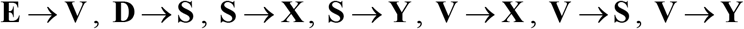

Above structural formulation implies that the observe expression profiles **X** is not a direct readout of the mechanism-relevant state **S** associated with drug combination **D**, and the observed outcome **Y** is not determined by tumor-intrinsic sensitivity alone. Instead, **X** and **Y** are jointly shaped by shared mechanism-related factors, unstable and non-transferable factors. As a result, although **S** denotes the transportable mechanism-relevant state, the observational relationship between **S** and **Y** may be distorted by host context, and other domain-specific influences. In causal terms, it induces back-door paths **S** ← **V** → **Y** between **S** and **Y**, such that the observational distribution cannot be directly interpreted as the net mechanism-related effect. Ideally, if the relevant dominant sources **V** of cross-domain heterogeneity are directly observed, one could block such back-door paths through standard adjustment. However, in the present setting, **S** and **V** are all latent and not directly observed, making it infeasible to specify a fully explicit adjustment set in the original observation space.

Therefore, we introduce two proxy latent representations: a causal latent representation *Z*_*S*_ and a domain-specific latent representation *Z*_*V*_,

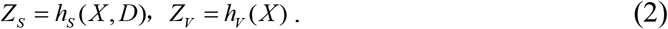

where *Z*_*S*_ is defined as a mechanism-aligned transportable surrogate representation, used to capture the shared, transportable mechanism-related information among cell lines and patients; *Z*_*V*_ is defined as a domain-specific response-modulating surrogate representation designed to absorb domain-specific heterogeneity and response-modulating variation, and serves as a proxy adjustment variable for latent back-door paths.

In the above representation view, the identification target of our framework is the interventional quantity

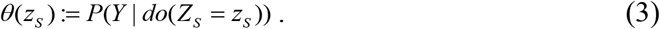

where, *P*(*Y* | *do*(*Z*_*S*_ = *z*_*S*_)) denotes the intervention distribution, *do*(·) denotes the ‘*do*’ operator ^63^. Notably, this identification target concerns the interventional definition quantity parameterized by *Z*_*S*_.

For the proxy-adjusted interventional estimand identification, we do not advocate strict back-door identification as the dominant sources of cross-domain confounding are not directly observed. Instead, we assume that *Z*_*V*_ provides a sufficient proxy for the main latent heterogeneity that jointly affects *Z*_*S*_ and *Y*. Under proxy ignorability, positivity, and consistency, the interventional quantity is approximately identified by the proxy-adjustment formula

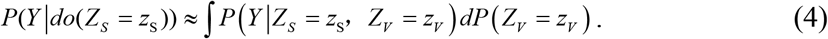

It should be interpreted as proxy-based approximate backdoor adjustment in the latent space, rather than as a strict classical backdoor identification with directly observed confounders. The proof is provided in Supplementary Information **Note** 3.

Separately from the identification of estimand, we interpret *Z*_*S*_ as a partially identifiable, mechanism-aligned surrogate representation, if it satisfies three properties: (i) it preserves the predictability of outcome **Y**; (ii) conditional on *Z*_*V*_, the relationship between *Z*_*S*_ and **Y** is approximately stable across domains; (iii) it carries the limited information of the domain label. The interpretation further relies on the assumption that the available environments are sufficiently heterogeneous to destabilize purely domain-specific spurious associations. In observation, there is no pseudo representation 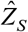 equivalent to *Z*_*S*_. Under these conditions, *Z*_*S*_ is better understood as capturing transportable mechanism-related information than as merely encoding stable statistical associations. The proof is provided in Supplementary Information **Note 4**.

Given that testing each drug combination in all possible individuals is impractical, and most drug combinations are often tested on a limited number of cell lines or patients, even some drug combinations do not have the testing samples, based on SCM above, we propose CaMeRe framework to predict the drug combination responses in clinical patients by jointly learning the representation *Z*_*S*_ and *Z*_*V*_.

### III. CaMeRe

CaMeRe is a bi-level domain generalization method based on meta-learning, which utilizes the advantages of sharing initialization parameters in meta-learning tasks to solve the problem of learning causal latent representation with few samples. In the limited number of labeled preclinical samples and a small set of unlabeled clinical samples, CaMeRe is used to predict the clinical patient response to drug combinations. In the following parts, we will introduce the core components in CaMeRe: Domain-Invariant Causal Representation Learning (**III.1**), Causally-Inspired Meta Representation Learning (**III.2**), and Inference and clinical test (**III.3**).

#### III.1 Domain-Invariant Causal Representation Learning

To learn domain invariant causal latent representations *Z*_*S*_ and domain-specific latent representation *Z*_*V*_, we reformulate the assumption conditions on *Z*_*S*_ and *Z*_*V*_ as information-theoretic object,

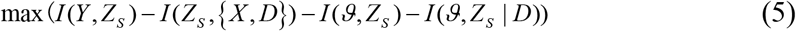

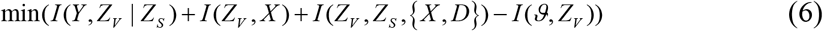

where, *I* (•) is mutual information, *ϑ* is domain label. Considering that the mutual information of high dimensional vectors is hard to estimate, we follow the rules of variational approximation to write *I* (*Y, Z*_*S*_) − *I* (*Z*_*S*_,{*X, D*}) as,

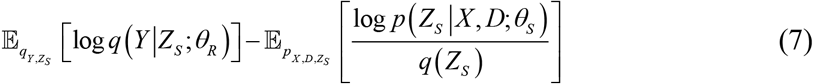

where, *q*(*Z*_*S*_) is the approximation to true marginal *p*(*Z*_*S*_), and *q* (*Y* | *Z*_*S*_;*θ*_*R*_) is the approximation to true marginal *p* (*Y* | *Z*_*S*_), *p* (*Z*_*S*_ | *X, D*;*θ*_*S*_) is the approximation to true posterior *p* (*Z*_*S*_ | *X, D*).

The mutual information terms *I* (*ϑ, Z*_*S*_) and *I* (*ϑ, Z*_*S*_ | *D*) can be expressed as 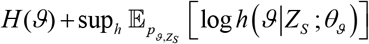, and 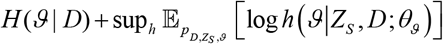. Here, sup denotes the least upper bound, *h* (*ϑ* | *Z*_*S*_;*θ*_*ϑ*_) is the approximation to true marginal *p* (*ϑ* | *Z*_*S*_), and *h* (*ϑ* | *Z*_*S*_, *D*;*θ*_*ϑ*_) is the approximation to true marginal *p* (*ϑ* | *Z*_*S*_, *D*), the entropy terms *H* (*ϑ*) And *H* (*ϑ* | *D*) are treated as constants. Similarly, we can transform *I* (*Z*_*V*_, *X*) − *I* (*ϑ, Z*_*V*_) as,

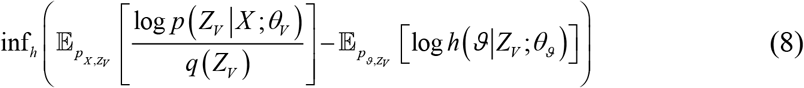

where, inf denotes the greatest lower bound, *q*(*Z*_*V*_) is the approximation to true marginal *p*(*Z*_*V*_), and *h* (*ϑ* | *Z*_*V*_;*θ*_*ϑ*_) to *p* (*ϑ* | *Z*_*V*_), *p* (*Z*_*V*_ | *X*;*θ*_*V*_) is the approximation to true posterior *p* (*Z*_*V*_ | *X*).

Rather than directly constraining *I* (*Y, Z*_*V*_ | *Z*_*S*_), which is challenging to estimate in high-dimensional latent space and does not directly represent the interventional effect of *Z*_*S*_ and *Y*. To better capture intervention invariant outcome mechanisms, we operationalize the minimization of *I* (*Y, Z*_*V*_ *Z*_*S*_) + *I* (*Z*_*V*_, *Z*_*S*_,{*X, D*}) using the following tractable objective,

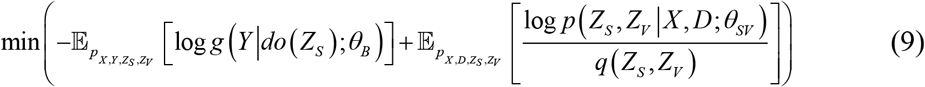

where, *q* (*Z*_*S*_, *Z*_*V*_) is the approximation to true marginal *p* (*Z*_*S*_, *Z*_*V*_); *p* (*Z*_*S*_, *Z*_*V*_ | *X, D*;*θ*_*SV*_) is the approximation to true posterior *p* (*Z*_*S*_, *Z*_*V*_ | *X, D*), and *g* (*Y* | *do* (*Z*_*S*_);*θ*_*B*_) is the approximation to true interventional distribution *p* (*Y* | *do* (*Z*_*S*_)).

For approximating above posterior distribution with deep learning, we parameterize the conditional likelihood using the neural networks. Specifically, we design a feature embedding module *f*_*E*_ (•;**Θ**_*E*_ = {**W**_*D*_,**W**_*S*_,**W**_*V*_,**b**_*S*_,**b**_*V*_}) composed of a drug feature encoder *f*_*ED*_ (•;**Θ**_*ED*_ = **W**_*D*_) to parameterize *p* ( *X*_*D*_ | *D*;*θ*_*ED*_), a causal feature encoder *f*_*ES*_ (•;**Θ**_*ES*_ ={**W**_*S*_,**b**_*S*_}), and a domain-specific feature encoder *f*_*EV*_ (•;**Θ**_*EV*_ ={**W**_*V*_,**b**_*V*_}) to parameterize *p* ( *X*_*S*_ | *X*;*θ*_*ES*_) and *p* ( *X*_*V*_ | *X*;*θ*_*EV*_); design a drug combination response predictor *f*_*R*_ (•;**Θ**_*R*_ ={**W**_*ø*_,**W**_*φ*_,**W**_*R*_,**b**_*ø*_,**b**_*φ*_,**b**_*R*_}) to parameterize *p* (*Z*_*S*_ | *X*_*S*_, *X*_*D*_;*θ*_*S*_) and *q* (*Y* | *Z*_*S*_;*θ*_*R*_), a unsupervised multi-domain discriminator *f*_*Dis*_ (•;**Θ**_*Dis*_ ={**W**_*ξ*_,**W**_*ζ*_,**b**_*ξ*_,**b**_*ζ*_}) to parameterize *h* (*ϑ* | *X*_*S*_;*θ*_*ϑ*_), *h* (*ϑ* | *X*_*S*_, *X*_*D*_;*θ*_*ϑ*_), and *h* (*ϑ* | *X*_*V*_;*θ*_*ϑ*_), and a back-door adjustment module *f*_*B*_ (•;**Θ**_*B*_ ={**W**_*τ*_,**W**_*ν*_,**W**_*B*_,**b**_*τ*_,**b**_*ν*_,**b**_*B*_}) to parameterize *p* (*Z*_*S*_, *Z*_*V*_ | *X*_*D*_, *X*_*S*_, *X*_*V*_;*θ*_*SV*_) and *g* (*Y* | *do* (*Z*_*S*_);*θ*_*B*_).

##### Feature Embedding Module

The feature embedding module *f*_*E*_ (•;**Θ**_*E*_) maps the raw inputs of each sample into the feature embedding space through the drug feature encoder, the causal feature encoder and the domain-specific feature encoder.

*For drug feature encoder*, each drug molecule is represented as an undirected graph *G*_*d*_ = (**V**_*d*_, **E**_*d*_, **X**_*d*_), where nodes **V**_*d*_ represent the atoms, edges **E**_*d*_ represent the chemical bonds connecting two atoms, and **X**_*d*_ are the node features extracted by encoding the atom-level descriptors such as atom type, degree, hybridization, and aromaticity, etc. We adopt a *L*_*d*_ layer message-passing-based graph neural network to extract the hierarchical molecular representations, which can flexibly capture structure and feature-level dependencies in the graph. At each layer *l*_*d*_, the representation 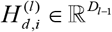 of each node *v*_*d,i*_ is updated by aggregating messages from its neighbors • (*v*_*d,i*_).

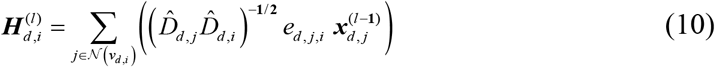

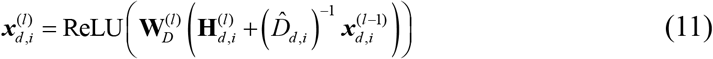

where, *e*_*d, j,I*_ represents the chemical bond link between *i*-th atom and *j*-th atom, 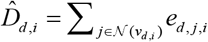 is the normalized degree of node 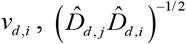 is the symmetric normalization term that is used to adjust the contribution of node for preventing highly connected nodes from dominating the feature propagation, 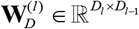 is a learnable transformation matrix, and ReLU(•) is a non-linear activation function.

After *L*_*d*_ message-passing layers, a readout function is applied to aggregate node-level representations into a fixed-dimensional molecular representation, and a multilayer perceptron (MLP) is used to map the graph-level representation into the desired output space.

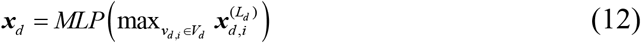

*For causal/domain-specific feature encoder*, it is used to map the gene expression profile 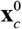 into the causal hidden representation 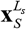 and domain-specific hidden representation 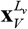. In this work, we adopt two multi-layer convolutional neural networks (CNNs) composed of *L*_*S*_ and *L*_*V*_ convolutional block, respectively. Each block applies a convolutional operation with kernel 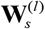 or 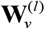, following by batch normalization *BN* (•) operator, a ReLU activation ReLU(•), and max-pooling operation *Pool* (•).

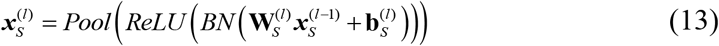

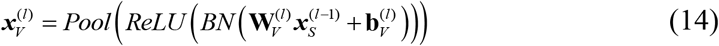

where, 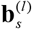 and 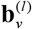 are the bias term of convolutional kernel in *l*-th convolutional block, respectively. After *L*_*S*_ or *L*_*V*_ convolutional blocks, these hidden representations are flattened and passed through a nonlinear transformation, typically a multilayer perceptron (MLP) with activation function (e.g., *ReLU*), for capturing the high-order features.

##### Drug Combination Response Predictor

In this module, drug combination features embedding ***x***_*D*_ = {***x***_*dA*_, ***x***_*dB*_} and the causal hidden representation ***x***_*S*_ are combined to parameterize the posterior distribution *p* ( ***Z***_*S*_ | ***x***_*S*_, ***x***_*D*_) of causal latent variable, from which the causal latent representation ***Z***_*S*_ is sampled by reparameterization trick^111^ and then used to predict drug combination response. Specifically, we model *p* ( ***Z***_*S*_ | ***x***_*S*_, ***x***_*D*_) as a diagonal multivariate Gaussian distribution *p* ( ***Z***_*S*_ | ***x***_*S*_, ***x***_*D*_)=*N* (***μ***_*S*_, diag((***σ***_*S*_)^2^). Here, the mean ***μ***_*S*_ and variance (***σ***_*S*_)^2^ are learned via two one-layer fully neural networks, respectively,

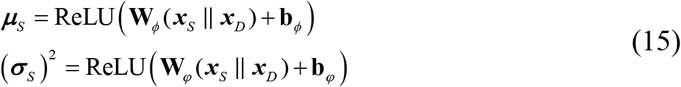

The drug combination-conditioned causal latent representation ***Z***_*S*_ taken from *p* ( ***Z***_*S*_ | ***x***_*S*_, ***x***_*D*_) can be represented as,

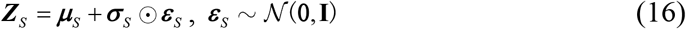

where ⨀ indicates the element-wise product.

For each sample, drug combination response predictor *f*_*R*_ can output its response value **ŷ**.

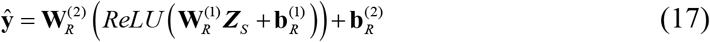

where, 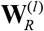 and 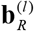 denote the weights and biases of *l*-th layer fully neural network, respectively.

To mitigate the pronounced class-imbalance between response and non-response samples, we adopt the balance SoftMax function ^112^ to map the logits output **ŷ** ^m^ of class *m* to conditional probability 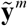,

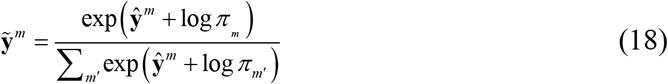

where *π* _*m*_ denotes the empirical prior of *m*-*th* class. In this work, *π*_*m*_ is the label frequency of the *m*-*th* class in a batch samples.

In this predictor, the sufficiency loss •_1_ is defined as follows:

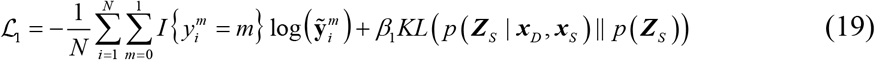

where, *N* is the total number of the labeled samples; 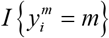 is the Kronecker indicator function, that is, 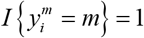 if the label of *y*_*i*_ belongs to *m*-*th* class, otherwise 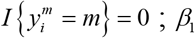 is a hyperparameter used to control the regularization strength.

##### Unsupervised multi-domain discriminator

Since explicit discrete domain labels are unavailable, we design an unsupervised multi-domain discriminator *f*_*Dis*_ to promote domain-invariant representations. Instead of predicting the domain label from a single point embedding, our multi-domain discriminator aims to evaluate the distinguishability of feature distributions learned in multiple domains for determining whether domain-invariant representations have been achieved, which is desirable for domain generalization. Specifically, given a batch of inputs, *f*_*Dis*_ first generates the high-order latent distribution *q* ( ***z*** | ***x***) for each input, and its distribution characteristics [***μ***^*ϑ*^; log(***σ*** ^*ϑ*^)^2^] are learned via two neural networks, respectively,

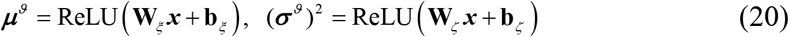

where, **W**_*ξ*_ and **W**_*ζ*_ are the weight matrixes of two networks, **b**_*ξ*_ and **b**_*ζ*_ are the bias, respectively.

For domain distribution of drug-conditioned causal latent representation 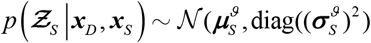, *f*_*Dis*_ takes the connection vector 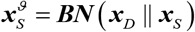 of drug combination feature embedding ***x***_*D*_ and causal hidden representation ***x***_*S*_ as input to output the mean 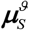 and variance 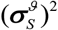. For domain distribution 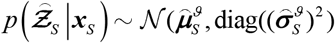 of drug-averaged causal latent representation, *f*_*Dis*_ takes the connection vector 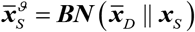 (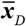 is the mean of multiple drug combination feature embeddings) as input to output the mean 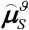 and variance 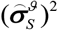. For domain distribution of domain-specific latent representation 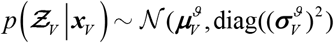,*f*_*Dis*_ takes domain-specific hidden representation *x*_*V*_ as input to output the mean 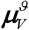 and variance 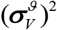. Then, *f*_*Dis*_ adopts Wasserstein distance 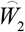 as a principle and geometry-aware metric to evaluate the discriminability between any two distributions. The calculation process of 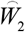 is detailed in Supplementary Information **Note 5**.

In this discriminator, we design following three loss functions: ℒ_2_, ℒ _3_ ℒ_4_, ℒ_2_ is a causal invariance loss used to ensure distribution invariance of causal latent representation associated with a specific drug combination across heterogeneous domains (*e.g*., cell lines and patients). ℒ_3_ is a domain independence loss used to ensure the distribution invariance of the causal latent representation on different cell lines and patients. ℒ_4_ is a domain dependence loss used to ensure that the distribution of domain-specific latent representations is distinguishable.

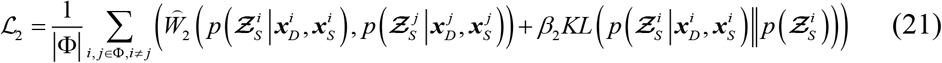

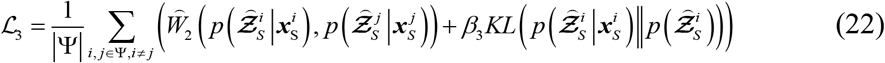

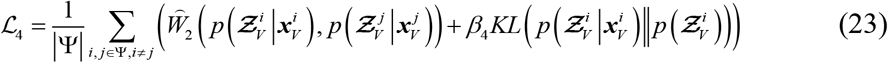

where, Φ denotes the set of sample domain distribution, Ψ denotes the set of individuals (*e.g*., cell lines and patients).

##### Proxy-Based Approximate Backdoor Adjustment module

Backdoor adjustment module *f*_*B*_ performs interventions on individual features by modifying individual domain-specific features, enabling the generation of counterfactual individual features and prediction of counterfactual individual combination response. Specifically, *f*_*B*_ first maps a batch of raw inputs (*i.e*., gene expression profiles) of individual patient/cell to high-level individual domain-specific representation space Ω. In space Ω, each individual patient/cell has a different domain feature representation.

Then, given a drug combination feature embedding ***x***_*D*_ and an individual causal hidden representation ***x***_*S*_, *f*_*B*_ adopts *do* operator to keep ***x***_*D*_ and ***x***_*S*_ unchanged, and assigns the individual domain-specific hidden representation ***x***_*V*_ to one of *K*_*V*_ individual domain-specific representations randomly selected from space Ω. The post-intervention distribution is defined as *p* ( ***Z***_*SV*_ | ***x***_*D*_, ***x*** _*S*_, ***x***_*V*_)=• (***μ*** _*SV*_, diag((***σ***_*SV*_)^2^)), where mean ***μ***_*SV*_ and variance (***σ***_*SV*_)^2^ are learned via two neural networks, respectively,

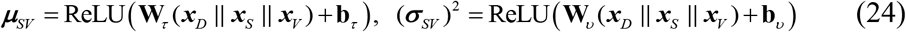

To assess the causal interventional invariance, we sample the joint latent representation ***Z***_*SV*_ from *p* ( ***Z***_*SV*_ ***x***_*D*_, ***x***_*S*_, ***x***_*V*_) using the reparameterization trick^111^, and then give the post-intervention individual combination response ŷ _*do*(*S*)_ predicted by a multi-layer fully connected neural network and *do* operator.

According to Eq. (18), we adopt the balanced SoftMax function to map the logit output 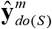 of class *m* to the conditional probability 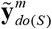, then define an intervention invariance loss ℒ_5_ to ensure the predicted drug combination response 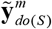 is not affected by domain-specific factors intervention.

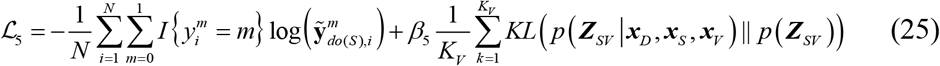

where, *N* is the total number of the labeled samples, *I* {·} is the Kronecker indicator function, *K*_*V*_ is the total number of interventions.

#### III.2 Causally-Inspired Meta Representation Learning

We introduce meta-training and meta-test phases in the causally-inspired meta representation learning procedure of CaMeRe, and realize meta-learning at tissue-specific task level. During meta-training phase, we adopt bi-level optimization strategy to train CaMeRe, aiming to achieve small sample bi-level domain generalization. For each task, our CaMeRe involves three processes: simulate multi-domain generalization, inner domain-invariant causal representation learning, and outer domain-invariant causal representation learning.

##### Simulate Multi-Domain Generalization

To simulate multi-domain generalization scenario from preclinical to clinical translation, we use tissue-specific cell line samples and patient samples to build support set and query set for each task. Specifically, for tissue T, we first build a support set 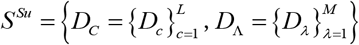 by randomly choosing *L* cell lines and *M* unlabeled patients from the tissue *T*, where each cell line *c* contains *K* samples 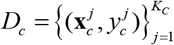 and each unlabeled patient contains one sample *D*_Λ_ = {**x**_*λ*_}. Due to the limited number of labeled patient samples in tissue *T*, which will increase the risk of overfitting during meta learning, we take the cell lines **C**′ as auxiliary domains, and then build a query set 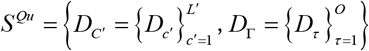 by randomly choosing *L*′ cell lines **C**′ ( **C** • **C**′ =Ø) and *O* labeled patients from the tissue *T*, where each cell line *c*′ contains 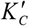 samples 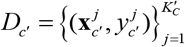, and each patient contains one sample *D* _*τ*_ = {(**x**_*τ*_, *y*_*τ*_)}.

To address the sparsity of treatment data caused by the limited or even absent cell lines (or patients) treated with a specific drug combination, we augment the support set 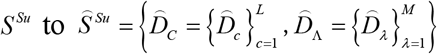 by assuming that every cell line and unlabeled patient in the support set have been treated with every drug combination in *S* ^*Su*^ set, and also augment the query set *T*^*Qu*^ to 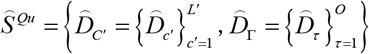. By the way, the augmented samples are unlabeled samples, which are used solely for unsupervised learning.

##### Inner/Outer-loop domain-invariant causal representation learning

We adopt the parameter grouping and multi-optimizer joint training strategy to perform inner-loop/outer-loop Domain-Invariant Causal Representation Learning (DICRL), for achieving finer grained, more stable, or more effective training. Let **Θ** = {**Θ**_*DS*_, **Θ**_*EV*_, **Θ**_*BR*_, **Θ**_*Dis*_} is the parameter set of inner-loop DICRL for each task, and 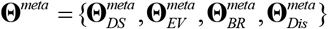 is the parameter set of outer-loop DICRL for all tasks. Here, **Θ**_*DS*_ = {**Θ**_*ED*_,**Θ**_*ES*_} is the parameter set in drug feature encoder *f*_*E*_ and causal feature encoder *f*_*ED*_, **Θ**_*EV*_ is the parameters in domain-specific feature encoder *f*_*E*V_, **Θ**_*BR*_ = {**Θ**_*B*_, **Θ**_*R*_} is the parameter set in Back-door Adjustment Module *f*_*B*_ and Drug Combination Response Predictor *f*_*R*_, **Θ**_*D*is_ is the parameters in Unsupervised Multi-Domain Discriminator *f*_*D*_. When DICRL is adapted to a new task, the parameters **Θ**^*meta*^ of outer-loop DICRL from all tasks will be updated into the parameters **Θ** by a few gradient descent updates on this new task. Especially, to enable adversarial optimization, we employ a gradient reversal layer (GRL) among the drug feature encoder, causal feature encoder and unsupervised multi-domain discriminator, for discovering which gradients from the adversarial loss are reversed before propagating to the feature embedding module. This setting encourages the modules to learn feature representations that are predictive for the main task but invariant to source and target distributions.

For each task with support set 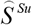, we use stochastic gradient descent to jointly train all blocks. At each iteration, CaMeRe performs inner-loop domain-invariant causal representation learning through the following steps,

1. update **Θ**_*Ds*_ to minimize sufficiency loss 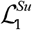 and intervention invariance loss 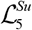, and maximize the causal invariance loss 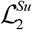 and domain independence loss 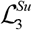,

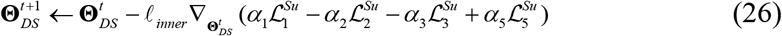
2. update **Θ**_***Ev***_ to minimize domain dependence loss 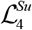 and intervention invariance loss 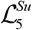,

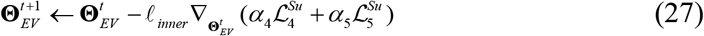
3. update **Θ**_*BR*_ to minimize sufficiency loss ℒ_1_ and intervention invariance loss ℒ_5_,

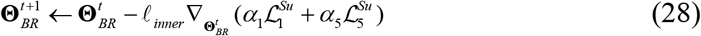
4. update **Θ**_*Dis*_ to minimize causal invariance loss 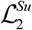, domain independence loss 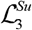, and domain dependence loss 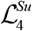,

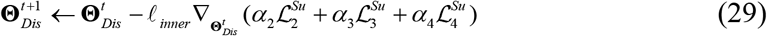

where, *ℓ*_*inner*_ is the learning rate of inner-loop DICRL, *α*_1_, …,*α*_5_ are weight factors, and 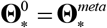.

For a batch of tasks with their query set 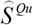, CaMeRe performs outer-loop DICRL by updating the shared initialization parameters 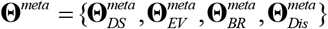 with four Adam optimizers based on the sufficiency loss 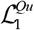, causal invariance loss 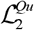, domain independence loss 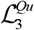, domain dependence loss 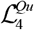, and intervention invariance loss 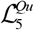

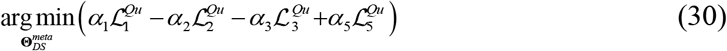

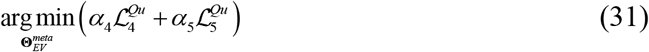

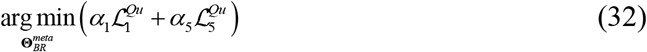

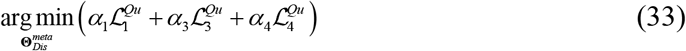

The above optimization operations encourage the learned causal latent representations to be domain-invariant, thereby facilitating generalization to the unlabeled target domain.

#### III.3 Inference and clinical test

In this module, we transfer the pretrained CaMeRe with multiple source tissues to new tissue *T*^*new*^, and update the parameters **Θ**^*meta*^ with support set 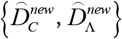 through inner DICRL process, aiming to adapt CaMeRe to new tissue *T*^*new*^. After that, given a clinical patient sample and a drug combination, CaMeRe can inference its causal latent representation, and then predict the patient clinical response to the drug combination.

### IV. Permutation SHAP

The precise calculation of Shapley values requires enumerating all subsets of features, which is intractable for high-dimensional models. Considering that our CaMeRe is a complex multi-head input and multi-head output architecture, the model-type-specific SHAP approximation approach (*e.g*., DeepSHAP) is not well-suited for providing reliable or effective explanations for CaMeRe, therefore, we employ the Permutation SHAP ^112-114^ (details provided in Supplementary Information **Note 6**) to attribute the model’s predictions to input features.

### V. Baseline Models

To comprehensively evaluate the effectiveness of CaMeRe, we selected several representative methods closely related to our work. Specifically, a traditional machine learning method (*i.e*., XGBoost) was included to assess performance in few-shot learning setting, two representative drug synergy prediction methods (*i.e*., DeepSynergy, MatchMaker) to evaluate domain generalization necessity, and six representative clinical drug response prediction methods (*i.e*., CODE-AE, ADAE, DSN-M/ DSN-A, TCRP, Velodrome) and to further demonstrate the advantages of CaMeRe over other domain generalization approaches. According to their core strategies for addressing domain shift, we grouped these methods into three categories: feature engineering-based methods, domain adaptation-based methods, and domain generalization-based methods. To ensure a fair comparison under a unified data setting, we made only the minimal modifications required to adapt each method to the clinical drug combination response prediction task, while preserving their core ideas. All methods were trained using the same labeled samples and unlabeled samples (if some baseline methods require the unlabeled samples) as CaMeRe. Details of the model adaptations and hyperparameter selection are provided in the Supplementary Information **Note 1**.

#### Computational Resource

All experiments run on Linux operating system with 24Gx4 NVIDIA RTX 3090 GPU, 40x2.4GHZ Intel Xeon CPU, and 500 GB RAM. Other computation parameters are given in **Tables S13-15** and **Notes 7-8** in Supplementary Information.

## Acknowledgements

We acknowledge the support of the National Natural Science Foundation of China (62473312 and 62173271 to S.W.Z., 62502163 to T.H.Z.).

## Author contributions

Q.Q.Z., S.W.Z, and T.H.Z. conceived and designed the study. Q.Q.Z. implemented CaMeRe. Q.Q.Z. performed the benchmarking studies with the assistance of M.H.S. Q.Q.Z. conducted the computational analysis with the assistance of J.N.L. and Y.R.Q. S.W.Z and T.H.Z mentored and guided the study. Q.Q.Z., S.W.Z, and T.H.Z. wrote the manuscript. All authors read and approved the final manuscript.

